# Targeted Epigenetic Modulation Outperforms Nuclease- and Deaminase-Based Editing for Durable *Pcsk9* Silencing in a Clinically Relevant Delivery System

**DOI:** 10.64898/2026.03.20.713290

**Authors:** Anusorn Mudla, Dominic Dimitri Quintana, Christian Flynn Atallah, Lindsey Renee Savoy, Angel I-Jou Leu, Thanhchau Dam, Grishma Acharya, Kumar Rajappan, Padmanabh Chivukula

## Abstract

Inhibition of proprotein convertase subtilisin/kexin type 9 (PCSK9) lowers low-density lipoprotein cholesterol, a major risk factor for cardiovascular disease. Although several gene therapy strategies targeting *Pcsk9* have been developed, direct comparisons across modalities are limited. To address this, we systematically evaluated cytosine base editing, nuclease-based CRISPR–Cas9, and epigenetic gene editing for *Pcsk9* suppression. We first engineered a cytosine base editor to introduce a premature stop codon, then optimized and characterized an epigenetic editor, and finally delivered all modalities as mRNA formulated in Arcturus lipid nanoparticles (LUNAR^®^) into wild-type mice, benchmarking them against conventional CRISPR–Cas9 and GalNAc-siRNA. Remarkably, epigenetic editing achieved the most efficient and sustained repression of PCSK9, maintaining low protein levels throughout the entire 30-day study period. By comparison, cytosine base editing reduced PCSK9 with minimal double-stranded DNA breaks and off-target effects, but editing precision requires further improvement, while GalNAc-siRNA produced only transient suppression, limiting its suitability for a one-time therapeutic approach. Collectively, these findings highlight the superior durability and efficacy of epigenetic gene editing and provide proof-of-concept for its combination with LUNAR^®^ delivery as a promising strategy for long-lasting hepatic-targeted therapy.

## Introduction

Secreted PCSK9 targets the low-density lipoprotein receptor (LDLR) for degradation in the liver. Following internalization of the LDLR–LDL complex, binding of extracellular PCSK9 redirects the receptor toward lysosomal degradation rather than recycling back to the cell surface ^1,2^. Consequently, reduction of PCSK9 increases LDLR recycling and surface expression, thereby enhancing hepatic clearance of circulating LDL particles and reducing cardiovascular risk ^3,4^. Individuals with homozygous loss-of-function mutations in *PCSK9* exhibit markedly lower LDL cholesterol levels without apparent adverse health effects^5–8^, providing compelling human genetic evidence supporting therapeutic inhibition of PCSK9. Monoclonal antibodies such as evolocumab^9^ and alirocumab^10^ inhibit PCSK9 function but require frequent dosing every 2–4 weeks and are typically used in combination with statin therapy^11^. More recently, inclisiran, a GalNAc-conjugated siRNA, has demonstrated durable LDL-C lowering with twice-yearly dosing^12–14^. While these approaches have shown clinical benefit, the development of a true “one-and-done” therapy motivates exploration of gene editing technologies capable of inducing long-lasting *PCSK9* suppression.

CRISPR–Cas systems have emerged as powerful tools for gene editing, including disruption of *Pcsk9* expression^15^. The programmability of single-guide RNAs (sgRNAs) enables efficient targeting of specific genomic loci^16–18^, and multiple Cas9 variants have been engineered to enhance specificity and reduce off-target activity, such as the high-fidelity SpCas9-HF1 variant designed to minimize non-specific DNA interactions^19,20^. Despite these advances, conventional CRISPR–Cas9 editing introduces double-stranded DNA breaks (DSBs), which are repaired through error-prone pathways that generate insertions or deletions^21^ and may lead to chromosomal rearrangements^21,22^. Although this mechanism is effective for gene disruption, it raises safety concerns, particularly for therapeutic applications. Moreover, many genetic diseases require precise nucleotide modifications rather than gene disruption alone. To address these limitations, base editing technologies were developed. By coupling a catalytically impaired Cas9 with cytosine or adenine deaminases, base editors enable targeted C•G-to-T•A or A•T-to-G•C conversions respectively without inducing DSBs^23–25^. While base editing has rapidly advanced toward clinical application^26^, challenges including limited editing efficiency, bystander mutations, and off-target activity remain to be addressed.

Given the inherent risks associated with permanent genome modification, epigenetic editing has emerged as a promising alternative strategy for gene therapy. Unlike genome editing, epigenetic editing does not alter the underlying DNA sequence but instead modulates gene expression by reshaping the epigenetic landscape^27,28^. Epigenetic editors typically consist of a DNA-binding domain fused to a transcriptional repressor, such as the Krüppel-associated box (KRAB) domain or DNA methyltransferases, enabling targeted and durable transcriptional repression ^29,30^. The DNA-binding component may be RNA-guided, as in CRISPR-based systems^31^, or protein-guided, as in zinc finger proteins (ZFPs)^27^ or transcription activator-like effectors (TALEs)^32,33^. Because epigenetic editing avoids DSBs and permanent sequence alterations, it substantially reduces risks associated with genomic instability and off-target mutagenesis. Importantly, epigenetic repression is potentially reversible through modification of effector domains, offering an additional layer of therapeutic control.

Efficient and targeted delivery remains a central challenge for gene therapy. For *Pcsk9* inhibition, hepatocytes, which constitute approximately 70% of liver cells^34^, represent the primary cell type of therapeutic relevance. Viral delivery systems such as adeno-associated virus (AAV) have demonstrated high transduction efficiency^35^ but are associated with safety concerns, including immunogenicity and the potential for genomic integration^36^. In contrast, recent advances in lipid nanoparticle (LNP)-mediated mRNA delivery offer a non-viral alternative with rapid development cycle, manufacturing scalability, improved safety, and transient expression profiles, well suited for gene editing applications^37,38^. Moreover, LNP composition can be tuned to enhance hepatocyte specificity and plasma clearance, enabling rapid formulation screening and improved therapeutic potency and safety^39,40^.

Although programmable epigenetic repression of *Pcsk9* has been demonstrated *in vivo*^41^, a direct comparison of a clinically relevant, LNP-delivered, mRNA-encoded epigenetic editing platform with advanced genome editing strategies, including high-fidelity CRISPR–Cas9, cytosine base editing, and GalNAc-conjugated siRNA, has not been previously reported. Here, we show that a single administration of a *Pcsk9*-targeted epigenetic editor achieves robust and sustained *Pcsk9* suppression, with efficacy exceeding that of genome editing and GalNAc-siRNA-based approaches.

## Results

### Development of a CRISPR cytosine base editor to introduce a premature stop codon in the *Pcsk9* open reading frame

Previous studies targeting *Pcsk9* have predominantly employed adenine base editors to disrupt mRNA splicing by modifying conserved sequences at exon–intron junctions, thereby introducing premature stop codons during translation^42^. To distinguish our approach, we pursued an alternative strategy to abrogate PCSK9 expression by directly introducing a premature nonsense mutation within the open reading frame using a cytosine base editor (CBE). Introduction of an early stop codon is expected to enhance gene disruption through protein truncation and increased nonsense-mediated mRNA decay^43,44^. Our CBE consists of a high-fidelity SpCas9 variant harboring the D10A nickase mutation (SpCas9-HF nickase), fused to the *Petromyzon marinus* cytidine deaminase (PmCDA1) and a *Bacillus subtilis* bacteriophage uracil–DNA glycosylase inhibitor (UGI) (Figure 1A). In this configuration, the Cas9 nickase introduces a single-strand nick in the non-complementary DNA strand^45^, enabling cytidine deamination within the editing window^46,47^. The UGI component prevents base excision repair of the resulting uracil ^23,48^, allowing its persistence through DNA replication and resulting in a stable C•G-to-T•A transition at the target locus, with a corresponding G•C-to-A• T mutation on the opposite strand.

**Figure 1.**
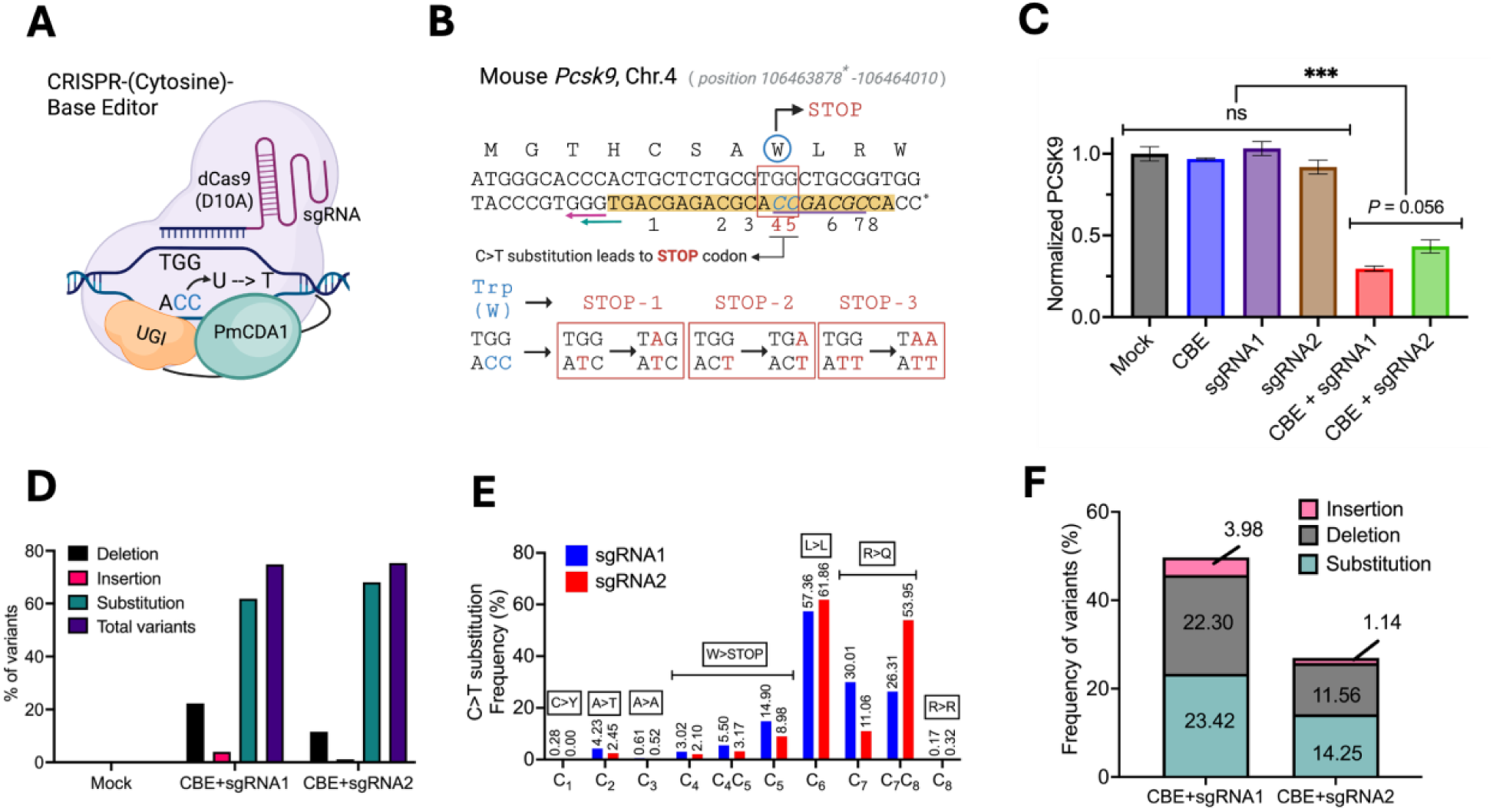
Development of cytosine base editors targeting mouse *Pcsk9* gene. (A) Schematic of the CRISPR cytosine base editor (CBE) complex composed of a high-fidelity SpCas9 nickase (SpCas9-HF), the cytidine deaminase PmCDA1, and a uracil–DNA glycosylase inhibitor (UGI) bound to the target DNA. Created with BioRender.com. (B) DNA sequence corresponding to the first 11 amino acids of the mouse *Pcsk9* open reading frame. C* denotes position 106,463,878 on mouse chromosome 4. Magenta and green arrows indicate the PAM sequences for sgRNA1 (GGG) and sgRNA2 (TGG), respectively. The 20-nucleotide sgRNA target region upstream of the PAM is highlighted in yellow. Numbers 1–8 indicate cytosine positions analyzed in panel E. The predicted CBE editing window (nucleotides 12–18 upstream of the PAM) is underlined and italicized. The red box highlights the tryptophan codon and the two cytosines whose C-to-T conversion generates three possible stop codons (shown in blue). (C) Secreted PCSK9 levels in culture media quantified by ELISA and normalized to mock (eGFP mRNA) transfection. Hepa1-6 cells were transfected with CBE mRNA and the indicated sgRNAs. Data were analyzed by one-way ANOVA followed by Tukey’s multiple-comparisons test (\*\*\**P* < 0.001). Error bars: SD. (D) Percentage of mutation variants identified by next-generation sequencing (NGS). “Total variants” represent the proportion of sequencing reads containing at least one mutation, with each read counted once regardless of the number of variant types present. (E) Frequency of C-to-T substitutions at each cytosine position within the sgRNA1 and sgRNA2 target sequences (positions indicated in panel B). Corresponding synonymous and nonsynonymous amino acid changes are indicated. (F) Frequency of variants generating premature stop codons. Substitution frequencies represent the combined C-to-T conversion rates at cytosine positions 4 and 5.

To guide sgRNA selection, we used BE-Designer from CRISPR RGEN Tools^49^. The coding sequence of exon 1 of the mouse *Pcsk9* gene, corresponding to amino acids 1–72, was provided as input, with the base editing type set to “BE (C to T)” and the editing window defined as positions 12–18. BE-Designer identified three candidate sgRNAs capable of generating premature stop codons. Two sgRNAs, sgRNA1 and sgRNA2, targeted tryptophan at position 8. Because tryptophan is encoded exclusively by the TGG codon, C-to-T conversion on the complementary strand generates TAA, TAG, or TGA stop codons on the coding strand (Figure 1B). The third sgRNA targeted glutamine at position 51 (CAG), for which C-to-T conversion produces only a single stop codon (TAG). We therefore prioritized the tryptophan-targeting sgRNAs, as they introduce an earlier premature stop codon and allow multiple possible C-to-T conversion outcomes that reliably generate translational termination.

We next compared the editing efficiency of sgRNA1 and sgRNA2 in Hepa1-6 mouse hepatocyte cells. CBE-gRNA-complex treatment reduced secreted PCSK9 levels by up to 60-70% (Figure 1C). Since sgRNA1 and sgRNA2 differ by only a single nucleotide within the 20-nucleotide protospacer sequence, no statistically significant difference in PCSK9 suppression was observed between the two guides. Nevertheless, in the absence of CBE or sgRNA complexes, no reduction in secreted PCSK9 was detected. To confirm on-target base editing, we performed next-generation sequencing (NGS) on cells transfected with eGFP mRNA (control) or CBE co-transfected with sgRNA1 or sgRNA2. Both sgRNAs produced comparable total mutation frequencies (74.82% and 75.37%, respectively), consistent with the observed reduction in PCSK9 protein levels. The majority of detected variants were base substitutions (>61%) rather than insertions or deletions (Figure 1D). Further analysis revealed that C-to-T substitutions predominated within the predicted editing window, occurring at frequencies of up to 61.86% across targeted cytosines (Figures 1B and 1E).

NGS analysis further demonstrated that CBE generated both synonymous and nonsynonymous mutations (Figure 1E and S1). Although C-to-T conversion at position C6 occurred at high frequencies (sgRNA1: 57.36% and sgRNA2: 61.86%), it did not alter the encoded leucine residue (Figures 1E and S1). Similarly, combined C-to-T conversions at positions C7–C8 occurred frequently (sgRNA1: 56.32% and sgRNA2: 65.01%) but resulted in an amino acid substitution (R10Q) with unknown effects on PCSK9 function (Figures 1E and S1). Importantly, C-to-T conversions at positions C4–C5, which generate premature stop codons, were detected at modest but meaningful frequencies (Figures 1B and 1E). The combined stop-codon-generating C-to-T frequency reached 23.42% for sgRNA1 and 14.25% for sgRNA2 (Figure 1E-F). Because 1-2 nucleotide insertion and deletion mutations can also introduce frameshifts and premature termination^50^, we considered the combined contribution of substitutions, insertions, and deletions when evaluating overall editing efficacy (Figure 1F). Based on these analyses, sgRNA1 was selected for all subsequent experiments.

### Stable suppression of *Pcsk9* transcription requires both DNMT and KRAB domains

Among epigenetic mechanisms of transcriptional repression, one of the best-characterized pathways involves two classes of regulators: Krüppel-associated box (KRAB)–containing zinc finger proteins and DNA methyltransferases (DNMTs)^51,52^. Previous studies have shown that epigenetic editors incorporating either KRAB or DNMT family proteins can repress reporter constructs and endogenous genes^27,31,53^. The catalytic activity of DNMT3A is known to be enhanced by its regulatory cofactor DNMT3L, and constructs containing C-terminally linked DNMT3A and DNMT3L exhibit greater gene silencing potency than DNMT3A alone^27,54–56^.

To assess the contribution of epigenetic effector domains to stable *Pcsk9* repression, we first characterized a previously published epigenetic gene editor (epi-editor)^27^ in cultured cells. Specifically, we asked whether a single epigenetic effector domain or a combination of effector domains is required to establish durable transcriptional repression. To this end, we generated three epi-editor variants composed of a DNA-binding domain (DBD) fused to either individual or combined effector domains: the catalytic domain of DNMT3A (cdDNMT3A), full-length DNMT3L, and the KRAB domain of ZNF10 (Figure 2A). All epi-editor variants shared the same DBD, consisting of an array of six engineered zinc finger proteins (ZFPs) targeting a locus within the mouse *Pcsk9* 5′ untranslated region (5′UTR) located in a CpG island (Figure 2B). Two control constructs were also generated: (1) a DNMT–KRAB fusion lacking the ZFP DBD (designated “(–) ZFP Ctrl”) to control for non-targeted effector activity, and (2) the ZFP DBD alone (“ZFP Ctrl”) to account for potential transcriptional interference due to steric hindrance. To enhance mRNA stability and translational efficiency, all constructs were codon-optimized using a proprietary algorithm.

**Figure 2.**
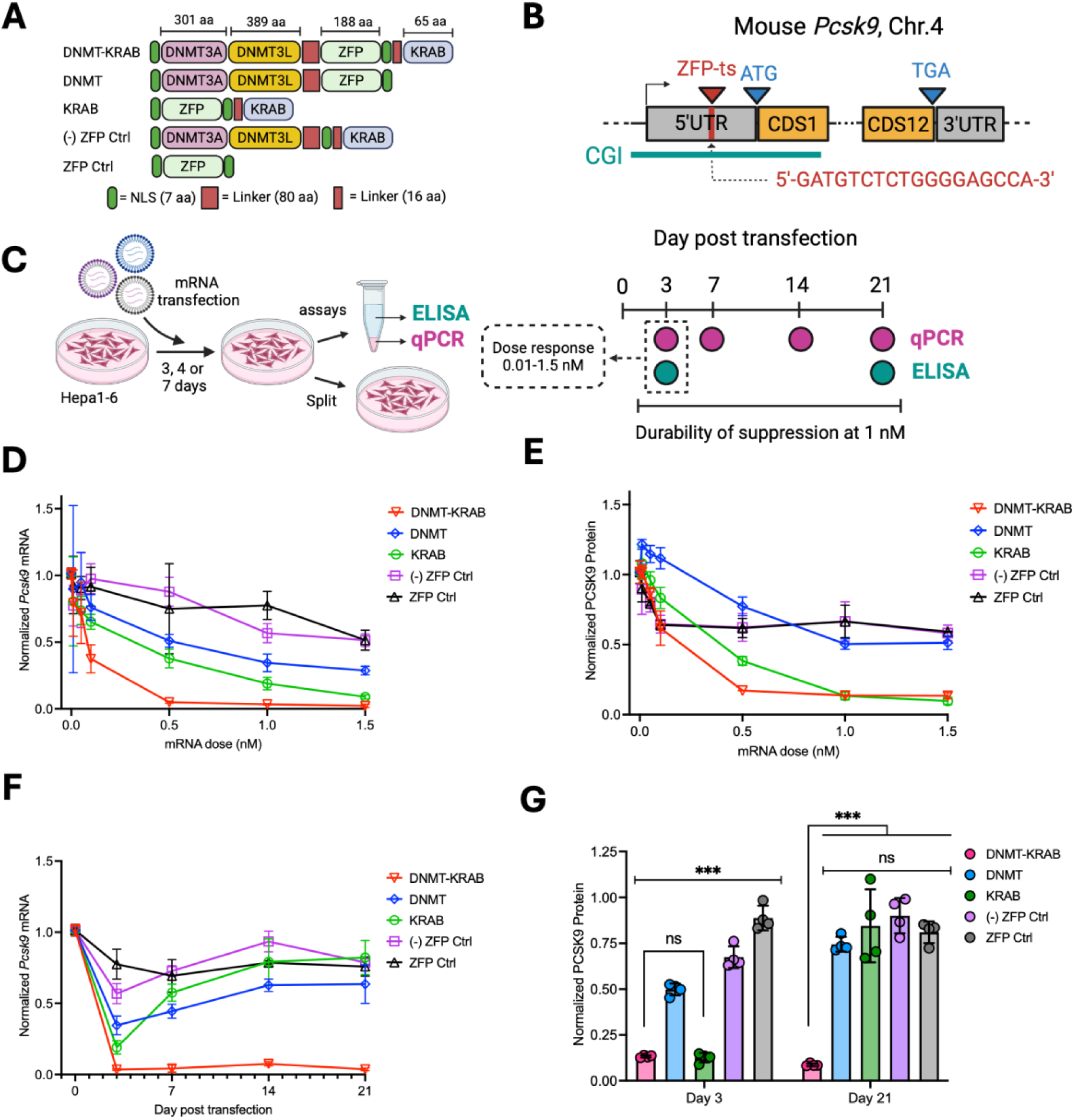
Epigenetic gene editors effectively suppress transcription *in vitro*. (A) Schematic representation of the molecular domains comprising the epigenetic gene editors and control constructs. The number of amino acids in each domain is indicated. (B) Schematic of the mouse *Pcsk9* gene showing the zinc-finger protein target site (ZFP-ts) and its sequence (in red) and the CpG island (CGI) region in green, with positions relative to the start codon (ATG) and stop codon (TGA). (C) Overview of the experimental workflow used to evaluate the efficacy of different epigenetic gene editor constructs and controls. Hepa1-6 cells were transfected with varying doses of mRNA using Lipofectamine™ MessengerMAX™. At the indicated time points, cells and culture media were collected for quantification of *Pcsk9* transcripts by RT–qPCR and secreted PCSK9 protein by ELISA. For the dose response, cells were transfected with mRNA at 0.01 – 1.5 nM and the samples were collected on day 3 post transfection. For the durability of suppression, cells were transfected at 1 nM and measured for *Pcsk9* transcript on day 3, 7, 14 and 21, and secreted PCKS9 protein was quantified on day 3 and day 21. Created with BioRender.com. (D–E) *Pcsk9* transcript levels measured by TaqMan RT–qPCR (D) and secreted PCSK9 protein levels measured by ELISA (E) on day 3 post-transfection with increasing mRNA doses (0.01–1.5 nM). Data were normalized to mock-transfected controls. (F) Line plot showing *Pcsk9* mRNA levels at the indicated time points following transfection with 1 nM mRNA, normalized to mock controls. (G) Bar plot showing secreted PCSK9 protein levels in the culture media on days 3 and 21 following transfection with 1 nM epigenetic gene editor mRNA. Error bars: SD. Statistical significance was determined by one-way ANOVA with Tukey’s multiple comparisons test (\*\*\**P* < 0.001), except where otherwise indicated.

We next performed dose–response studies to evaluate the potency of each epi-editor in Hepa1-6 cells. mRNA was delivered using Lipofectamine™ MessengerMax™ at concentrations ranging from 0.01 to 1.5 nM, followed by quantification of *Pcsk9* transcripts by TaqMan RT–qPCR and measurement of secreted PCSK9 protein on day 3 post-transfection (Figure 2C). All epi-editor variants significantly repressed *Pcsk9* transcription relative to mock (eGFP mRNA), (–) ZFP, and ZFP-only controls (Figure 2D). Notably, the dual-effector epi-editor (DNMT–KRAB) exhibited the greatest potency, with an IC₅₀ of 0.09 nM, compared with epi-editors containing single effector domains (DNMT: 0.39 nM; KRAB: 0.28 nM), representing up to a fourfold improvement in potency (Figure S2A). A similar trend was observed for secreted PCSK9 protein levels, with the DNMT–KRAB epi-editor achieving up to a sevenfold lower IC₅₀ relative to single-domain epi-editors (Figures 2E and S2B). Because transcriptional repression plateaued at a dose of 1 nM, this concentration was selected for subsequent durability studies.

To evaluate the stability of *Pcsk9* repression, Hepa1-6 cells were transfected with 1 nM of epi-editors or control constructs, and transcript levels were measured at days 3, 7, 14, and 21 post-transfection (Figure 2C). Strikingly, only the dual-effector DNMT–KRAB epi-editor maintained robust repression of *Pcsk9* transcription throughout the entire time course, whereas epi-editors containing DNMT or KRAB alone failed to sustain long-term silencing (Figure 2F). These results were consistent with PCSK9 protein measurements at days 3 and 21 (Figure 2G), indicating a strong correlation between transcriptional and protein-level suppression. Importantly, DNMT–KRAB reduced PCSK9 expression to as low as 8.7% of mock control levels at day 21. In contrast, firefly luciferase reporter analysis showed that protein expression from delivered mRNA declined by day 3 post-transfection (Figure S2C), strongly suggesting that the sustained repression mediated by DNMT–KRAB reflects stable epigenetic modification of the chromosomal locus rather than prolonged mRNA or protein persistence.

### Stable suppression of *Pcsk9* transcription requires temporal coordination of KRAB and DNMT effector domains

Closer examination of the dose–response data revealed that at high concentrations, a single KRAB-based epi-editor achieved *Pcsk9* suppression comparable to that of dual-effector constructs (Figures 2D-E). However, neither KRAB nor DNMT alone was sufficient to sustain transcriptional repression over time (Figures 2F-G). In contrast, simultaneous presence of both effector domains, as in the DNMT-KRAB epi-editor, resulted in durable gene silencing.

DNMT3A/3L mediates DNA methylation, a well-established mechanism of transcriptional repression, whereas the KRAB domain recruits histone-modifying enzymes to establish repressive chromatin states^57^ (Figure 3A). Given these distinct yet complementary mechanisms, we asked whether sequential delivery of the two effector domains could also induce stable transcriptional repression. To answer this, Hepa1-6 cells were first transfected with 1 nM DNMT or KRAB epi-editor mRNA using Lipofectamine™ MessengerMax™. DNMT-KRAB and eGFP mRNA served as positive and mock controls, respectively. After 24 hours, cells were washed to remove residual mRNA and subsequently re-transfected with the alternate effector domain, eGFP, or left untreated. Cell lysates were collected on days 7, 14, and 21 following the initial transfection (Figure 3B).

**Figure 3.**
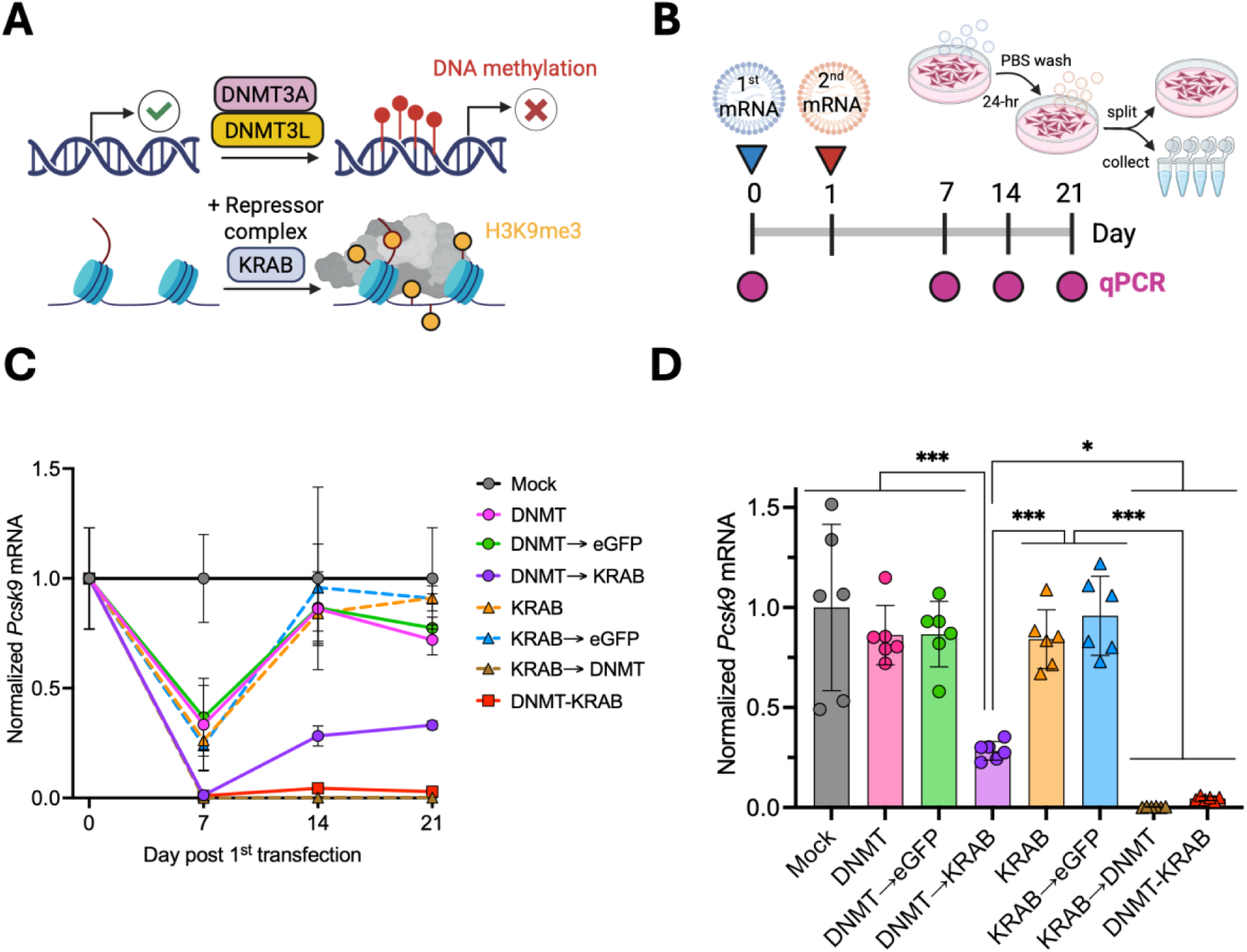
Stable transcriptional repression of *Pcsk9* requires coordinated activity of DNMT and KRAB effector domains. (A) Schematic illustration of the distinct epigenetic mechanisms mediated by DNA methyltransferases (DNMTs; top), which establish DNA methylation, and the KRAB domain (bottom), which recruits histone-modifying complexes to induce repressive chromatin states. (B) Schematic of the sequential transfection experimental design. Hepa1-6 cells were transfected with 1 nM of the first mRNA, washed with PBS 24 hours post-transfection to remove residual mRNA, and subsequently transfected with 1 nM of the second mRNA. Cells were harvested at the indicated time points, lysed, and stored at −80°C for TaqMan RT-qPCR analysis, with a subset of cells retained for later time points. Both panel A and B were created with BioRender.com. (C) Time-course analysis of *Pcsk9* mRNA levels quantified by TaqMan RT-qPCR following sequential (mRNA1→ mRNA2) or single effector domain delivery. “DNMT-KRAB” is dual-effector control. Data were normalized to the mock (eGFP mRNA) transfection control. (D) Normalized *Pcsk9* mRNA levels at day 21 post-transfection. Error bars: SD. Statistical significance was determined by one-way ANOVA followed by Tukey’s multiple comparisons test (\**P* < 0.05, \*\*\**P* < 0.001).

Consistent with prior observations, epi-editors containing a single effector domain (DNMT or KRAB) failed to maintain long-term repression of *Pcsk9* transcription (Figure 3C). Transcript levels were reduced at day 7 but returned to the level similar to the mock control by day 14. In contrast, sequential delivery of DNMT and KRAB (DNMT → KRAB or KRAB → DNMT) resulted in *Pcsk9* silencing comparable to the DNMT-KRAB control at day 7. Notably, the KRAB → DNMT sequence maintained sustained repression through day 21, whereas the DNMT → KRAB sequence exhibited gradual transcriptional recovery after day 7, although expression levels remained significantly lower than those in mock-treated or single-effector groups (Figure 3D). Importantly, replacing the second effector with eGFP mRNA did not reproduce this repression, indicating that the observed effects were not attributable to transfection -related toxicity. Together, these results demonstrate that stable transcriptional repression can be achieved through sequential delivery of KRAB and DNMT domains within a 24-hour window.

The temporal coordination between DNA methylation and histone modification is apparently critical for establishing durable repressive chromatin states. The partial loss of repression observed in the DNMT → KRAB sequence suggests that DNA methylation alone, in the absence of timely histone modification, may not be stably maintained. This finding implies that the interval between DNA methylation and histone remodeling must be shorter than 24 hours, potentially reflecting cell-cycle–dependent regulation^58,59^. Conversely, the sustained repression observed in the KRAB → DNMT sequence suggests that histone-based repression is more persistent and may better tolerate cell-cycle progression prior to DNA methylation^60,61^. While defining the precise temporal requirements of these processes will require further experimentation, our findings highlight the importance of effector timing in epi-editor–mediated gene regulation and provide mechanistic insight into the temporal coordination underlying stable epigenetic silencing.

### Epigenetic gene editing provides superior repression of *Pcsk9* transcription

CRISPR–Cas9 is widely regarded as a benchmark nuclease-based gene-editing modality. Accordingly, we directly compared the *Pcsk9* knockdown efficiency of the dual-domain epi-editor with that of CRISPR–Cas9 (gene editor) and a cytosine base editor (base editor) (Figure 4A). Throughout this section, the epi-editor refers to the dual-effector epigenetic editor (DNMT-KRAB) described above. For the gene editor, we used the SpCas9-HF variant, which is identical to the Cas9 backbone used in the base editor but lacks the D10A mutation. Chemically modified sgRNAs used for the gene and base editors targeted the same genomic locus, which differed from the target site of the epi-editor (Figure 4B).

**Figure 4.**
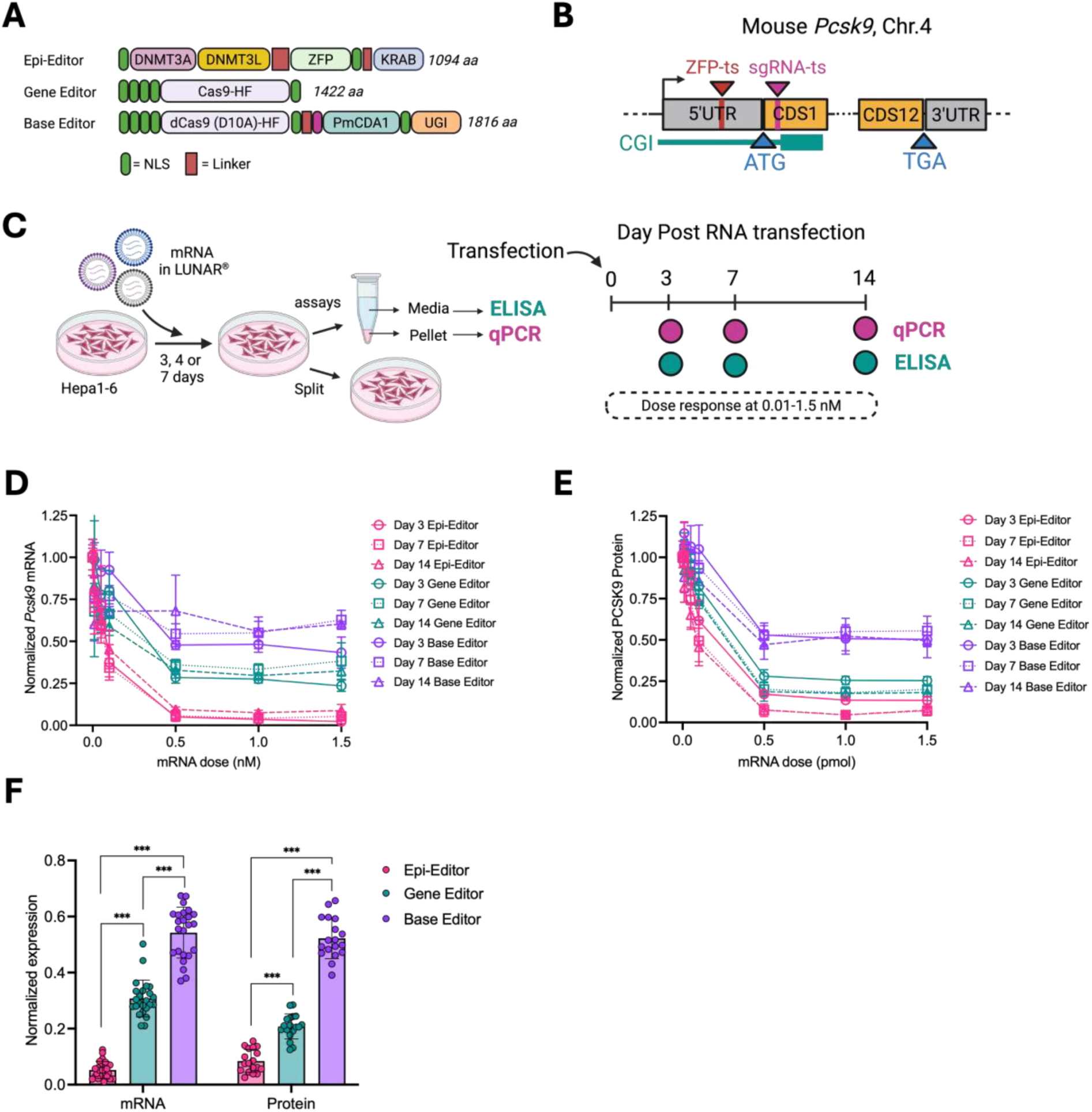
Comparative effect of *Pcsk9* suppression by distinct gene-editing modalities *in vitro*. (A) The domain architectures of the epigenetic gene editor (epi-editor), CRISPR–Cas9–based gene editor (gene editor), and cytosine base editor (base editor). The open reading frame (ORF) size for each modality is indicated. (B) *Pcsk9* locus showing ZFP target site for epi-editor (ZFP-ts) and gRNA target site (sgRNA-ts) for gene/base editors. (C) Experimental workflow for comparing the efficacy of the different gene-editing modalities. Hepa1-6 cells were transfected with varying doses of mRNA using LUNAR®. At the indicated time points, cell lysates and culture media were collected for TaqMan RT-qPCR analysis of *Pcsk9* transcripts and ELISA quantification of secreted PCSK9 protein, respectively. Created with BioRender.com. (D–E) Dose- and time-dependent suppression of *Pcsk9* mRNA transcripts (D) and secreted PCSK9 protein (E). Data were normalized to the mock (eGFP mRNA) transfection control. (F) Comparison of *Pcsk9* mRNA and PCSK9 protein reduction pooled from Hepa1-6 cells transfected with 1 and 1.5 nM mRNA across days 3, 7, and 14. Statistical significance was determined by one-way ANOVA followed by Tukey’s multiple comparisons test (\*\*\**P* < 0.001). Error bars: SD.

We next performed dose–response experiments in Hepa1-6 cells by transfecting mRNA encoding the epi-editor, or mRNA and gRNA at a 1:2 molar ratio for the gene- and base-editors. *Pcsk9* transcript levels in cell lysates and secreted PCSK9 protein levels in culture media were quantified on days 3, 7, and 14 post-transfection (Figure 4C). Across all doses tested, the epi-editor consistently outperformed both gene and base editors in suppressing *Pcsk9* expression (Figures 4D and 4E). Suppression began to plateau at approximately 0.5 nM. At maximal effect, the epi-editor reduced *Pcsk9* transcript levels to 5.3% of mock control, whereas the gene and base editors achieved reductions of only 29.7% and 53.6%, respectively. Transcriptional repression by all three modalities persisted through day 21 (Figure S3A). Although the calculated IC₅₀ values were comparable (epi-editor, 0.077 nM; gene editor, 0.107 nM; base editor, 0.066 nM), these values alone do not adequately reflect relative potency because the modalities did not achieve equivalent maximal suppression (Figure S3B).

To further compare the suppression effect across timepoints, we analyzed transcriptional and protein suppression at combined doses of 1 and 1.5 nM across multiple time points. The overall suppression consistently ranked as epi-editor > gene editor > base editor for both *Pcsk9* mRNA and PCSK9 protein levels (Figure 4F). As expected, transcriptional and translational repression were strongly correlated in the epi-editor group, reflecting direct epigenetic silencing of gene transcription. In contrast, the gene editor group exhibited a significant divergence between mRNA and protein suppression (70.3% versus 79.2%, *P* < 0.001) (Figure S3D), suggesting that indels generated by double-strand breaks did not substantially reduce *Pcsk9* transcription but instead impaired translation of full-length protein. Based on similar reasoning, a comparable effect was anticipated for the base editor. However, mRNA and protein suppression in this group were well correlated (46.4% versus 47.8%, *P* = 0.902) (Figure S3D), indicating that base substitutions may influence transcriptional regulation differently from indel-based mutations. Elucidating the underlying mechanisms of this distinction warrants further investigation but lies beyond the scope of the present study. Collectively, these results demonstrate that the epi-editor is a more effective transcriptional repressor of *Pcsk9* than either CRISPR–Cas9 or cytosine base editing approaches in this experimental setting.

### LNP-mediated delivery of gene therapy modalities results in persistent *Pcsk9* silencing in mice

Effects observed *in vitro* often demonstrate limited translatability to *in vivo* settings due to multiple factors, including constraints associated with the delivery system. Although LNP have demonstrated robust delivery efficiency, challenges remain with respect to cell type targeting and potential cellular toxicity. To validate the *in vitro* observations and to assess both the potency and safety of the gene therapy modalities, we evaluated the *Pcsk9* transcriptional suppressor approaches in mice. RNA cargos (mRNA alone or mRNA co-formulated with sgRNA) were encapsulated using Arcturus Therapeutics’ proprietary ionizable lipid. A GalNAc-conjugated siRNA targeting *Pcsk9* was included as a non-LNP delivery control. LNP-formulated eGFP mRNA served as a mock control to confirm that any observed reduction in plasma PCSK9 was not attributable to nonspecific LNP-associated toxicity, while a PBS-treated group was included as a safety control. Blood samples were collected via the retro-orbital plexus on days 0 (pre-dose), 3, 7, 14, 21, and 30 (Figure 5A). All animals were euthanized on day 30, and livers were harvested for clinical assessment. All RNA-loaded LNPs and GalNAc-siRNA formulations were well tolerated across treatment groups, and no animals required early termination. Body weights changes remained similar across all groups, including the controls (Figure S4A). Serum aspartate aminotransferase (AST) and alanine aminotransferase (ALT) levels were comparable to those of the PBS control group at all time points, indicating the absence of overt hepatotoxicity ^62^ and supporting the safety of the LUNAR^®^ formulation (Figures 5B-C).

**Figure 5.**
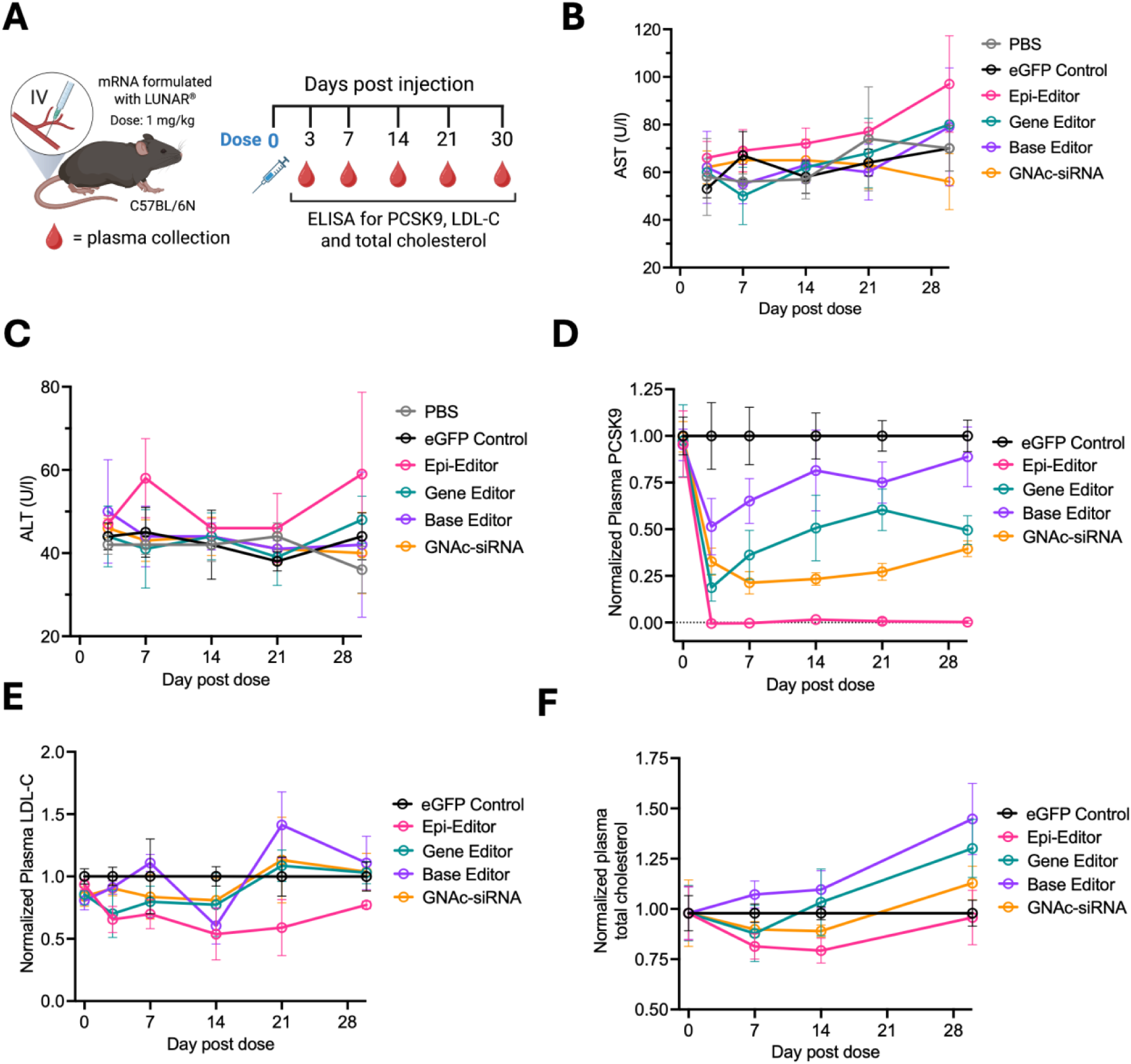
Epigenetic gene editor achieves durable PCSK9 suppression *in vivo*. (A) Experimental workflow. Hepa1-6 mRNA (epi-editor, gene editor, or base editor) and sgRNA were formulated LUNAR^®^ for systemic delivery, while GalNAc-conjugated siRNA was administered directly. eGFP mRNA served as a vehicle control. Blood samples were collected at the indicated time points for downstream analyses. Created with BioRender.com. (B–C) Serum aspartate aminotransferase (AST) and alanine aminotransferase (ALT) levels measured at each time point to assess hepatotoxicity. (D) Plasma PCSK9 protein levels quantified by ELISA. (E–F) Circulating low-density lipoprotein cholesterol (LDL-C) and total cholesterol levels. All plasma data (D–F) are normalized to eGFP control at the corresponding time point. Statistical analyses were performed as described in the Methods section. Error bars: SD.

A near-complete reduction in plasma PCSK9 was observed in the epi-editor–treated group (Figures 5D and S4B). When gene therapy modalities were compared at day 3, the time point associated with maximal PCSK9 suppression, the magnitude of reduction ranked as follows: epi-editor (100%), gene editor (81.3%), and base editor (48.6%). The GalNAc-conjugated siRNA control achieved its maximal reduction of 78.7% at day 7, with suppression remaining relatively stable (72.8–78.7%) for approximately two weeks before gradually declining to 60.5% by day 30. In contrast, PCSK9 suppression mediated by the epi-editor was sustained throughout the entire 30-day study period, indicating durable epigenetic modification (Figures 5D and S4B and S4E) and supporting efficient hepatocyte delivery of the LUNAR^®^ formulation. PCSK9 reduction mediated by both the gene editor and base editor declined more rapidly than that observed with the siRNA control. Notably, the base editor exhibited minimal residual activity by day 30, with only an 11.1% reduction relative to the eGFP control group (Figure S4E). Collectively, these results demonstrate superior and sustained repression of PCSK9 expression by the epigenetic editing modality compared with the other therapeutic approaches evaluated.

Circulating PCSK9 plays a central role in cholesterol homeostasis by binding to the low-density lipoprotein receptor (LDLR) on hepatocytes and promoting its lysosomal degradation ^63,64^. Consequently, reductions in plasma PCSK9 are expected to enhance LDLR recycling and lower circulating cholesterol levels. Despite the substantial suppression of PCSK9 achieved by the gene therapy modalities, particularly the epi-editor, no statistically significant reductions in plasma LDL-C or total cholesterol were observed across treatment groups (Figures 5E–F and S4C–D). The use of wild-type animals in this study may have contributed to the modest lipid response, as this model exhibits inherently low baseline LDL-C levels^65^ and limited dynamic range for further cholesterol lowering. The epi-editor group consistently exhibited lower LDL-C levels relative to controls and other modalities, although these differences did not reach statistical significance at most time points. The maximal LDL-C reduction in the epi-editor group was observed on day 14 (46.3%, *P* = 0.009), but this effect diminished by day 30 (22.8% reduction, *P* = 0.093). Unexpectedly, total cholesterol levels increased by an average of 19.7% at day 30 in the gene editor, base editor, and siRNA treatment groups (Figure S4E). In contrast, the epi-editor group exhibited a maximal total cholesterol reduction of 29.7% on day 14 (*P* < 0.001), consistent with the observed LDL-C trend; however, by day 30, the reduction decreased to 15.1% and was no longer statistically significant relative to controls (Figure S4E). These findings highlight an important limitation of wild-type models for evaluating lipid-lowering outcomes and suggest that metabolically sensitized or hyperlipidemic models may be required to fully capture the therapeutic impact of sustained PCSK9 repression on cholesterol metabolism.

## Discussion

The main objective of this study was to compare the efficacy of different gene therapy modalities for *Pcsk9* suppression within the same experimental framework. We evaluated a newly developed cytosine base editor, a traditional CRISPR–Cas9 gene editor, and a dual-domain epigenetic gene editor (epi-editor). Our results consistently demonstrated the superiority of the epi-editor over the other gene therapy modalities and the FDA-approved GalNAc-conjugated siRNA. Delivered as mRNA comprising a ZFP DNA-binding domain and two effector domains via the LUNAR^®^ formulation, the epi-editor achieved near-complete plasma PCSK9 silencing in mice for 30 days. In contrast, the gene and base editors, while significantly reducing PCSK9 levels, failed to sustain suppression, likely due to the outgrowth of unedited hepatocytes. This interpretation is consistent with our *in vitro* data, in which neither the gene editor nor the base editor achieved complete *Pcsk9* suppression even at the highest dose tested (Figure 4D), indicating comparatively lower editing efficiency. The sustained repression observed with the epi-editor supports stable inheritance of the induced epigenetic state and suggests highly efficient hepatocyte delivery mediated by the LUNAR^®^ system.

We acknowledge several limitations in our study. First, the cytosine base editor tested here may not represent the most optimized version. Previous studies using adenine base editors (ABE) achieved up to 90% PCSK9 reduction in macaques^42,66^, and such a system might have served as a stronger comparator to the epi-editor. Our choice of CBE was intended to expand understanding of this technology. Given constraints on producing a premature stop codon, we limited the target to the first exon of *Pcsk9* and tested only two sgRNAs suggested by the BE-Designer tool. As expected, CBE exhibited bystander C-to-T conversions^67^, confirmed by high-frequency unintended edits in NGS data (Figure 1E). Alternative engineered cytidine deaminases with narrower editing windows may improve specificity and reduce off-target effects^68^. Furthermore, the relatively high frequency of indels generated by CBE raises safety concerns that require additional investigation. Similarly, the gene editor was limited by sgRNA choice. Nonetheless, the combination of sgRNA1 and Cas9 -HF achieved ∼50% PCSK9 reduction at day 30, comparable to prior studies^27^, even at half the dose. Higher doses in previous reports ^27,31^ increased efficacy, suggesting that our *in vivo* base and gene editor dosing was suboptimal and explaining the discrepancy between persistent *in vitro* suppression at high doses and partial *in vivo* reduction after a single administration (Figures 4D–E and 5D). Moreover, high doses of CRISPR-based therapies present both formulation and toxicity challenges, emphasizing the advantages of epi-editors, which require only a single, shorter mRNA and lower doses. In addition, the epi-editor strategy described here eliminates the need for single-guide RNA–mediated targeting, distinguishing it from prior approaches that rely on sgRNA for locus specificity^31^. This reduction in molecular complexity streamlines *in vivo* delivery and mitigates challenges associated with sgRNA stability and co-delivery, further supporting the translational feasibility of this modality.

Another limitation is the choice of animal model. Despite robust PCSK9 suppression, statistically significant reductions in LDL-C and total cholesterol were not observed. Wild-type C57BL/6N mice on a normal diet have low baseline LDL-C because most circulating cholesterol is carried in HDL^65,69^. Even *Pcsk9* knockout mice show only modest changes in total plasma cholesterol despite substantial LDLR upregulation^70,71^. Additionally, hepatic LDLR levels in mice are already high, so further reduction of PCSK9 does not substantially increase LDL-C uptake^63,72^. Feedback mechanisms, including increased hepatic cholesterol synthesis, altered VLDL secretion, and changes in intestinal cholesterol absorption, may also limit cholesterol reduction ^72,73^. Future studies employing high-fat diet models, humanized apolipoprotein B (APOB) or LDLR mice, or hypercholesterolemic backgrounds may better reveal the physiological impact of PCSK9 inhibition^74–77^. In addition, studies in nonhuman primates whose lipoprotein metabolism more closely resembles that of humans will provide a highly relevant translational model for evaluating the therapeutic impact of PCSK9-targeting strategies^42^.

While mechanistic exploration of epigenetic state establishment was not the primary focus, we investigated how the temporal sequence of effector domain delivery influences silencing. Stable repression of *Pcsk9* required both DNMT and KRAB domains. Sequential delivery revealed that KRAB followed by DNMT produced durable silencing, whereas DNMT followed by KRAB resulted in gradual recovery. KRAB likely initiates transient chromatin repression by recruiting histone-modifying complexes, such as histone deacetylases (HDACs), histone methyltransferases (HMTs), and heterochromatin protein 1 (HP1), creating a permissive environment for DNMT-mediated DNA methylation^57^. Conversely, DNMT alone can transiently suppress transcription^59^, but without timely KRAB activity, long-lasting epigenetic marks are not established. Defining the duration of transient repression for individual effector domains could inform the design of optimized epi-editors.

Efficient delivery to target cells is critical for therapeutic efficacy. While our epi-editor design was adapted from Cappelluti et al.^27^, we codon-optimized the mRNA for improved expression and employed the LUNAR^®^ delivery system, resulting in higher efficacy and an improved safety profile. Although both studies had comparable designs (Figure S5A), the Arcturus epi-editor consistently outperformed the previously reported EvoETR-8, achieving 99.8% versus 75.9% PCSK9 reduction at day 30 with a three-fold lower dose (1 mg/kg vs 3 mg/kg) (Figures S5B–C). Both studies showed only modest effects on LDL-C and total cholesterol, reflecting limitations of the wild-type mouse model (Figures S5D–E). Importantly, LNP formulation largely determines safety^78–80^. The LNP used in Cappelluti et al. (“LNP-D”) produced markedly elevated AST and ALT levels, even three days post-dosing, with values 10.6- and 8.9-fold higher than baseline, respectively (Figures S6A-D). In contrast, LUNAR^®^ maintained low AST and ALT levels, and even at 3 mg/kg, transient enzyme elevations at 6 hours returned to baseline within two days (Figures S6E–F). These findings highlight the improved hepatotoxicity profile^81^ of LUNAR^®^, supporting its potency and tolerability. Future studies will investigate the cellular uptake, cell type distribution, and mechanisms underlying endosomal release.

In summary, codon-optimized epi-editor delivered via LUNAR^®^ demonstrated superior *Pcsk9* suppression compared with cytosine base and CRISPR–Cas9 gene editors. By avoiding double-stranded breaks and minimizing off-target mutagenesis, epi-editors offer a safer alternative to nuclease-based therapies. This study adds further evidence supporting mRNA-based epigenetic gene editing as a potentially low - cost, one- and-done therapeutic approach for hypercholesterolemia and other atherosclerotic cardiovascular diseases.

## Materials and Methods

### Construct design and RNA production

Fusion proteins were generated using codon-optimized mouse catalytic domains of DNA methyltransferase 3A (DNMT3A; GenBank: 13435), full-length DNA methyltransferase 3L (DNMT3L; GenBank: 54427), and the Krüppel-associated box (KRAB) domain of ZNF10 (GenBank: 7556). For CRISPR-based gene editing, a codon-optimized high-fidelity variant of Streptococcus pyogenes Cas9 (SpCas9-HF) containing four point mutations (N497A, R661A, Q695A, and Q926A) was used to minimize off-target cleavage. The cytosine base editor comprised codon-optimized SpCas9-HF carrying the D10A nickase mutation, the cytidine deaminase from Petromyzon marinus (PmCDA1), and a bacteriophage-derived uracil–DNA glycosylase inhibitor from Bacillus subtilis. All constructs were cloned into in-house–designed plasmids optimized for mRNA production. Chemically modified single-guide RNAs were synthesized by Integrated DNA Technologies. Each construct contained both N-terminal and C-terminal nuclear localization signals (PKKKRKV), and all plasmids were verified by Sanger sequencing.

Capped and polyadenylated mRNAs incorporating N1-methyl-pseudouridine were generated by in vitro transcription (IVT) using linearized plasmid templates and T7 RNA polymerase. IVT reactions were performed as previously described^38,82,83^, with proprietary modifications to enable efficient incorporation of a Cap 1 analog and to ensure high RNA quality. Transcribed mRNAs were purified using silica-based columns and quantified by UV absorbance. RNA integrity and size distribution were assessed by fragment analysis using an Advanced Analytical platform. Purified RNAs were resuspended in RNase-free water and stored at −80 °C until use.

### Next generation sequencing

Hepa1-6 cells were seeded at a density of 3.0 × 10⁵ cells per well in 6-well plates and transfected with 2 μg of mRNA encoding the base editor and 1 μM sgRNA. Twenty-four hours post-transfection, cells were harvested by gentle scraping, transferred to 15-mL conical tubes, and centrifuged at 1,000 × g for 5 min. Cell pellets were resuspended in 1 mL PBS, re-pelleted in 1.5-mL microcentrifuge tubes, and submitted to GeneGoCell (San Diego, CA) for library preparation and sequencing. Amplicon libraries were generated using GeneGoCell’s proprietary G-Amp^SM^ platform for library preparation with the following primers: forward, GCATCAGCTCTTCATAATCTCCA; reverse, GAGCCAAGTGCCCCGAGTC. Libraries were sequenced on an Illumina NextSeq 2000 platform. Sequencing data were analyzed using GeneGoCell’s G-Amp^SM^ bioinformatics pipeline to quantify editing outcomes and variant frequencies.

### mRNA-LNP formulation

Lipid nanoparticles (LNPs) encapsulating mRNA test articles were formulated by rapid microfluidic mixing of an ethanolic lipid solution with an aqueous RNA solution, as previously described^84^. Briefly, proprietary ionizable amino lipid, proprietary helper lipid, cholesterol (Avanti Polar Lipids), and DMG-PEG (1,2-Dimyristoyl-sn-glycerol methoxypolyethylene glycol, PEG chain molecular weight: 2000, NOF America Corporation) are dissolved in ethanol. An aqueous solution of the mRNA is prepared in citrate buffer (pH 3.5). The lipid mixture is then combined with the RNA solution at a flow rate ratio of 1:3 (V/V) using the NanoAssemblr^TM^ microfluidic system (Precision NanoSystems, Vancouver, Canada), resulting in spontaneous formation of LNPs. The mixed material was then diluted three times with HEPES buffer pH 8 after leaving the micromixer outlet which further reduced the ethanol content. The diluted LNP slurry was purified by tangential flow filtration with hollow fiber membranes (mPES Kros membranes, Repligen), and A total of 10 diavolumes were exchanged, effectively removing the ethanol. Concentration of the formulation is adjusted to the final target RNA concentration using 100,000 MWCO Amicon Ultra centrifuge tubes (Millipore Sigma) followed by filtration through a 0.2 µm PES sterilizing-grade filter. Particle size was determined by dynamic light scattering (ZEN3600, Malvern Instruments). Encapsulation efficiency was calculated by determining unencapsulated RNA content by measuring the fluorescence upon the addition of RiboGreen (Molecular Probes) to the LNP slurry (Fi) and comparing this value to the total RNA content that is obtained upon lysis of the LNPs by 1% Triton X-100 (Ft), where % encapsulation = (Ft − Fi)/Ft × 100. The final formulation was stored at −70 ± 10 °C until use.

### Cell culture and *in vitro* assays

Hepa1-6 mouse hepatocyte cells were obtained from ATCC and maintained in Dulbecco’s Modified Eagle’s Medium (DMEM; HyClone, Logan, UT) supplemented with 10% fetal bovine serum (FBS; HyClone) and 1% antibiotic–antimycotic (Thermo Fisher Scientific, Waltham, MA). Cells were cultured at 37 °C in a humidified incubator with 5% CO₂. For long-term durability studies, cells were passaged every 3–4 days at a 1:8 split ratio. Cells were seeded either in 96-well plates at 8,000 cells per well or in 24-well plates at 80,000 cells per well, depending on the assay format. For in vitro transfection experiments, cells were transfected with mRNA complexed with Lipofectamine MessengerMAX (Thermo Fisher Scientific) at final mRNA concentrations ranging from 0.01 to 1.5 nM. For CRISPR-based gene editor and cytosine base editor conditions, sgRNA was included during complex formation at an mRNA:sgRNA mass ratio of 1:2. Lipid–RNA complexes were prepared according to the manufacturer’s instructions.

Cell culture media were collected at designated time points, stored at −80 °C, and analyzed using a commercial pre-spotted ELISA kit (R&D Systems, Minneapolis, MN) following the manufacturer’s protocol. Cells were lysed using *Cells-to-CT™* cell lysis reagents (Thermo Fisher Scientific), stored at −80 °C, and analyzed by quantitative PCR using a QuantStudio instrument (Life Technologies, Carlsbad, CA). qPCR was performed using the TaqMan RNA-to-CT 1-Step Kit (Thermo Fisher Scientific) in a 384-well plate format with technical duplicates. Mouse *Pcsk9* and *Gapdh* TaqMan probes (Thermo Fisher Scientific) were used for gene expression analysis. Relative expression levels were calculated using the ΔΔCt method, normalized first to the housekeeping gene (*Gapdh*) and then to the experimental control. Pooled samples across all treatment groups were used to calibrate qPCR assay parameters. For luciferase assays of the reporter construct, cells were washed with PBS and lysed using 1× passive lysis buffer. Luminescence was measured using the Luciferase Assay System (Promega, E4550) according to the manufacturer’s instructions and quantified on a Cytation 3 plate reader (BioTek).

### Mouse handling and treatments

All animal procedures were approved by the Explora BioLabs Institutional Animal Care and Use Committee (IACUC) and conducted under Animal Care and Use Protocol EB-17-004-003. Female C57BL/6N mice (8 weeks of age; The Jackson Laboratory, Bar Harbor, ME) were used for all in vivo studies. Animals were randomly assigned to 10 experimental groups (n = 4 per group) and housed at the Arcturus Therapeutics vivarium under a 12-h light/12-h dark cycle with ad libitum access to food and water.

For in vivo administration, RNA–LNP formulations were dosed at 1 mg/kg in a 10mL/kg dose volume and administered via intravenous tail-vein injection. Blood samples corresponding to approximately 5–7% of total circulating volume were collected from the retro-orbital plexus using non-heparinized micro-hematocrit capillary tubes (Kimble Chase, Vineland, NJ) on days 0 (pre-dose), 3, 7, 14, 21, and 30. Blood was transferred to either EDTA-coated collection tubes or serum separator tubes, and centrifuged at 2,000 × g for 10 min at 4 °C. Plasma and serum were isolated respectively, aliquoted into single-use samples, and stored at −80 °C until analysis. At the study endpoint (day 30), mice were deeply anesthetized and euthanized by cervical dislocation.

### Blood analysis

Circulating PCSK9 levels in mouse plasma were quantified using a commercial pre-spotted ELISA kit (R&D Systems, Minneapolis, MN) according to the manufacturer’s instructions. Plasma samples were diluted 1:100 in assay diluent immediately prior to loading onto the ELISA plate. Plasma low-density lipoprotein cholesterol (LDL-C) was measured using a colorimetric assay kit (Novus Biologicals, Centennial, CO) following the manufacturer’s protocol, with samples loaded undiluted. Total cholesterol levels were quantified using a fluorometric assay kit (Cayman Chemical, Ann Arbor, MI), with plasma samples diluted 1:300 in sample diluent prior to analysis. Serum alanine aminotransferase (ALT) and aspartate aminotransferase (AST) levels were measured using a clinical chemistry analyzer (Vet Axcel; Alfa Wassermann, West Caldwell, NJ) according to the manufacturer’s standard operating procedures.

### Statistical analyses

Unless otherwise indicated, data are presented as mean ± standard deviation (SD). All statistical analyses and data visualization were performed using GraphPad Prism (GraphPad Software, San Diego, CA). Comparisons among groups were conducted using one-way analysis of variance (ANOVA), as appropriate for the experimental design. When a significant overall ANOVA effect was detected, group differences were assessed using planned multiple-comparison analyses, as specified in the Results section. Tukey’s multiple-comparisons test was applied for pairwise comparisons among all groups, assuming approximately Gaussian data distributions. Half-maximal inhibitory concentrations (IC₅₀) were determined using three-parameter logistic nonlinear regression models. Goodness of fit was evaluated using the coefficient of determination (R²) and residual standard deviation of regression (RSDR), as appropriate. All P-values are reported in the Results section, and values of P < 0.05 were considered statistically significant.

## Data Availability Statement

All data is available on reasonable request to the corresponding authors.

## Author Contributions

A.M., D.D.Q. designed research. A.M., D.D.Q. analyzed data and wrote manuscript. A.M., D.D.Q., C.F.A., L.R.S., A.I.L., and T.D. performed research. G.A., K.R., and P.C. supervised research.

---

**Figure S1.**
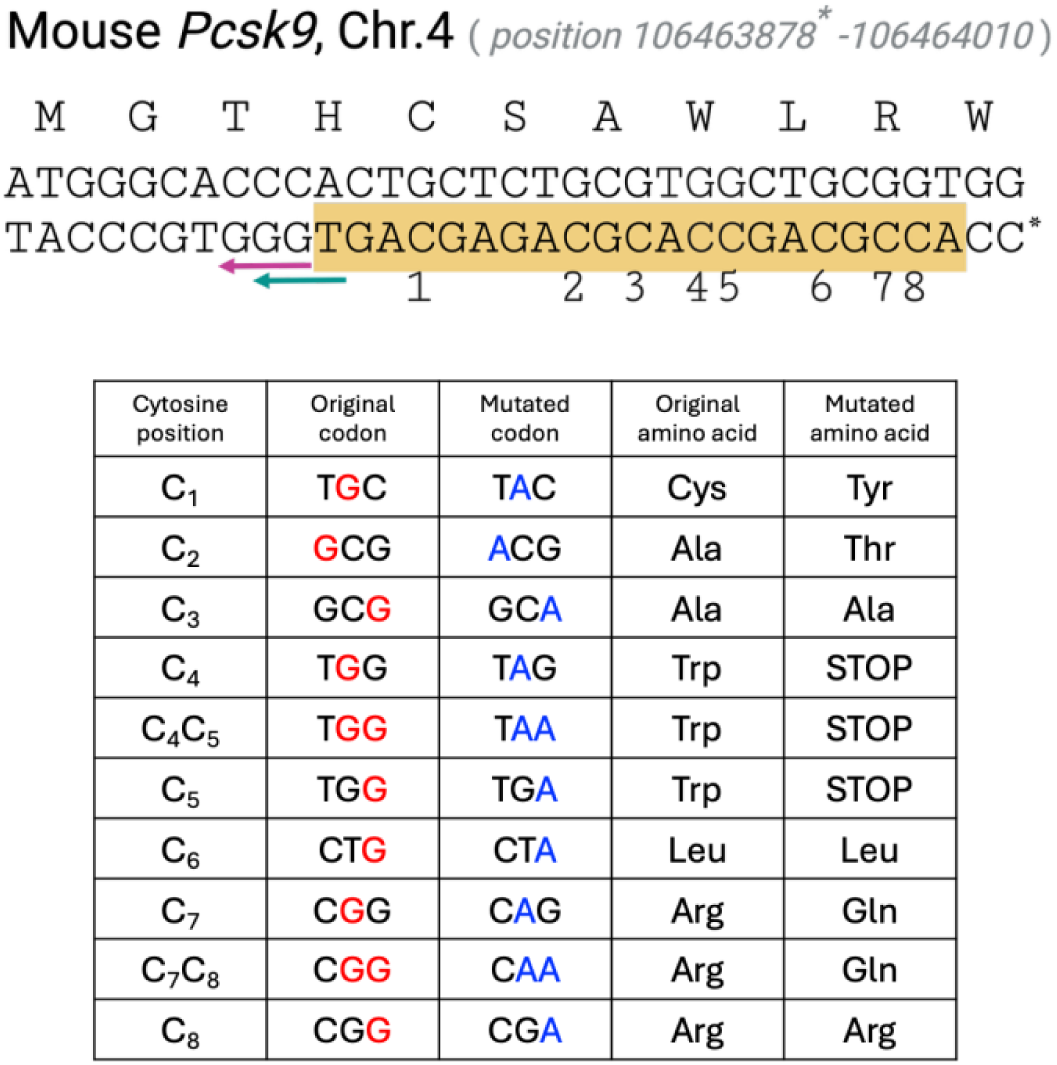
C-to-T conversion on the noncoding strand results in synonymous and nonsynonymous mutations in the *Pcsk9* gene. (Top) The nucleotide sequences of the coding (sense) and noncoding (antisense) strands corresponding to the first 11 amino acids of the mouse *Pcsk9* gene are shown. Magenta and green arrows indicate the PAM sequences for sgRNA1 (GGG) and sgRNA2 (TGG), respectively. C* denotes position 106,463,878 on mouse chromosome 4. The 20-nucleotide sgRNA target region upstream of the PAM is highlighted in yellow. Numbers 1–8 indicate positions of potential C-to-T conversions within the sgRNA-targeted region by the cytosine base editor. (Bottom) An overview of the predicted consequences of individual C-to-T conversions on the noncoding strand, resulting in corresponding G-to-A substitutions on the coding strand and the associated synonymous or nonsynonymous amino acid changes.

**Figure S2.**
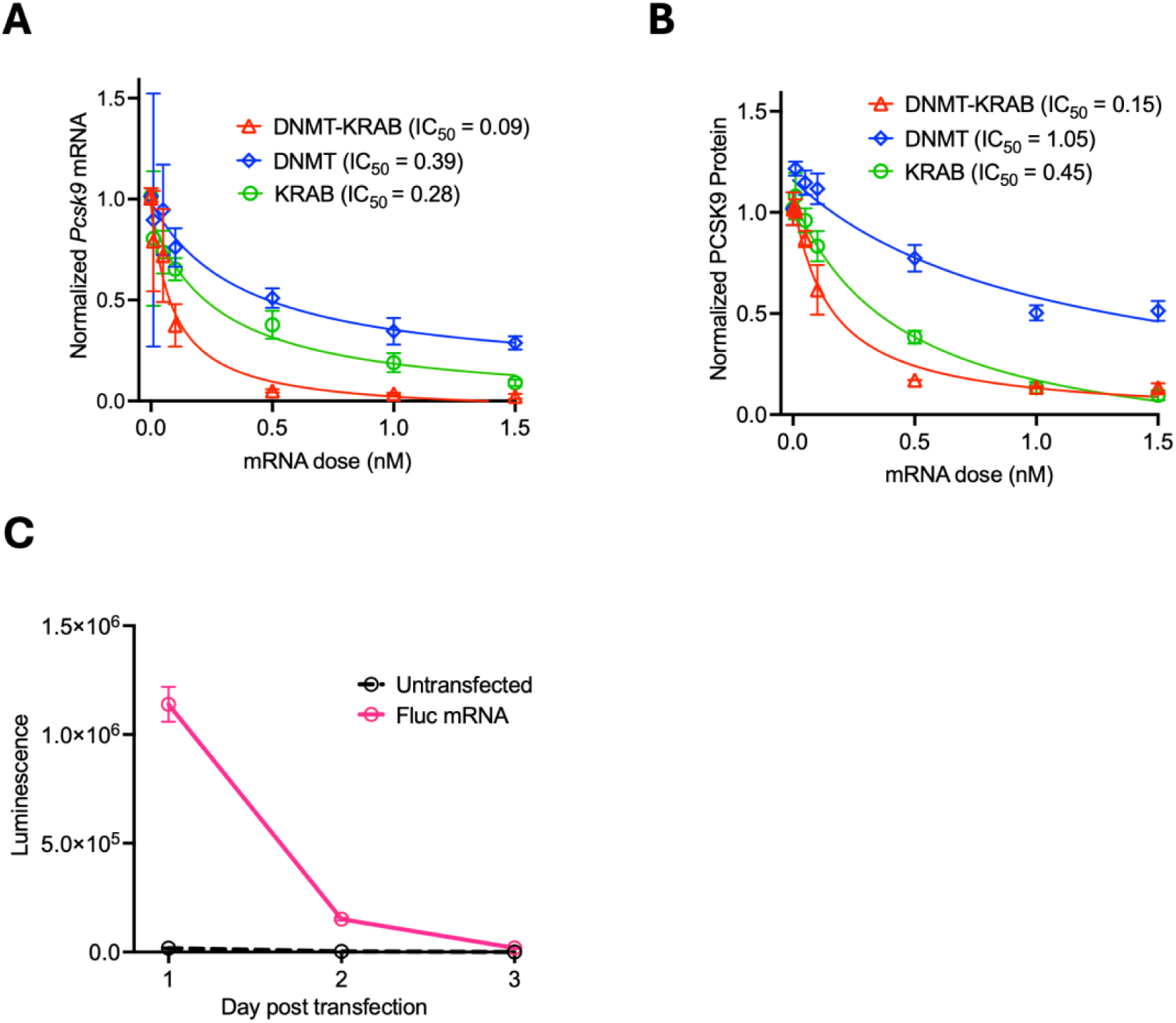
Dual epigenetic effector domains exhibit greater potency than single effector domains. (A–B) Nonlinear regression analysis of *Pcsk9* suppression measured by TaqMan RT–qPCR (A) and secreted PCSK9 protein quantified by ELISA (B) following transfection with epigenetic gene editor mRNAs. IC₅₀ determined using a three-parameter logistic regression model. (C) Firefly luciferase (Fluc) reporter expression in Hepa1-6 cells on days 1-3 following transfection with 1 nM mRNA. The reporter construct contains the same 5′ UTR and 3′ UTR as the epigenetic gene editor constructs. Error bars: SD.

**Figure S3.**
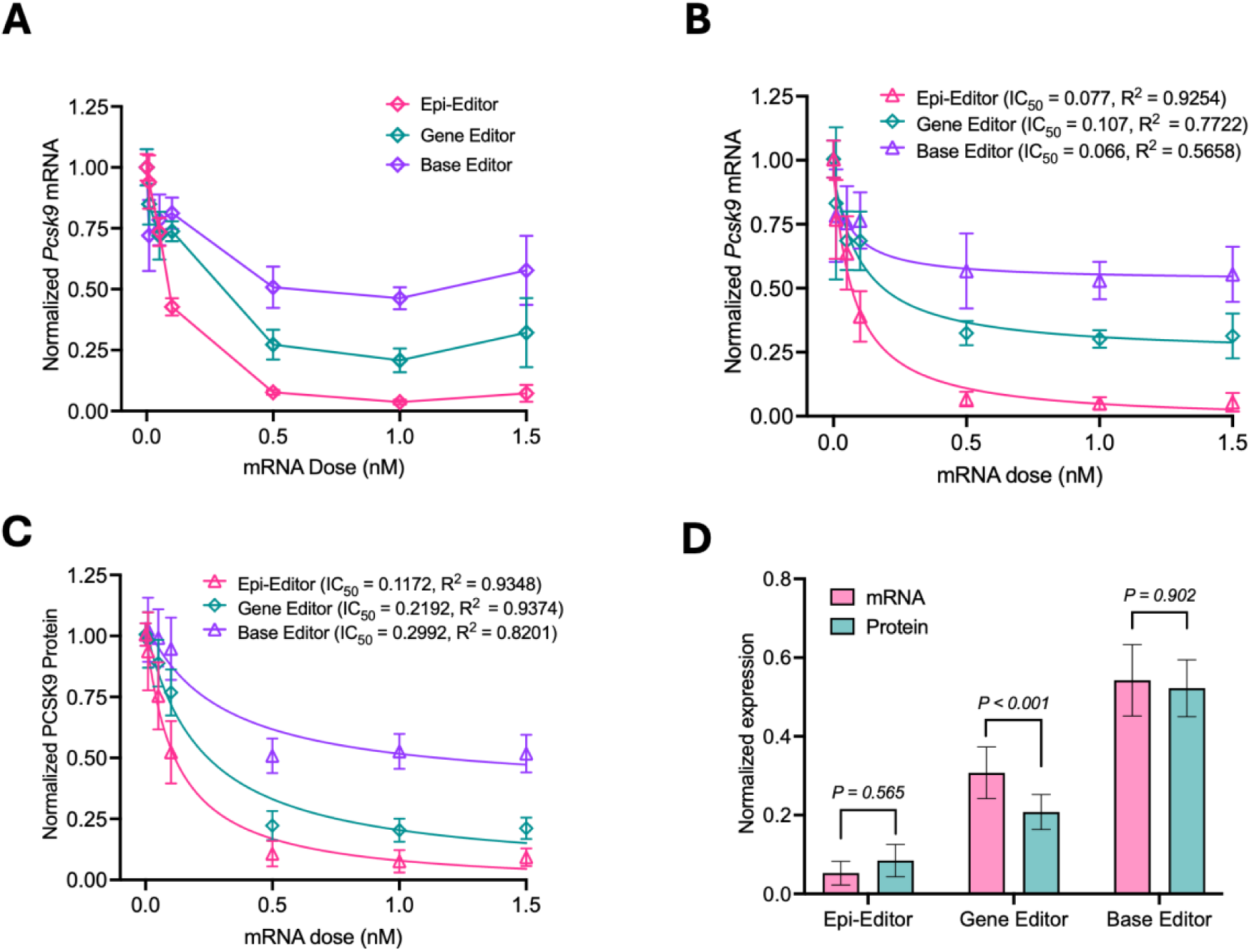
Epigenetic gene editor provides long-lasting and superior suppression of *Pcsk9* compared with gene and base editors. (A) Dose-dependent suppression of *Pcsk9* mRNA measured by TaqMan RT-qPCR on day 21 post-transfection in Hepa1-6 cells. Data were normalized to the mock control. (B–C) Nonlinear regression fits of *Pcsk9* mRNA (B) and secreted PCSK9 protein in culture media (C) following transfection with varying doses of epi-editor, gene editor, or base editor mRNAs. TaqMan RT-qPCR and ELISA were performed on days 3, 7, and 14. Data were normalized to the corresponding mock (eGFP mRNA) control, and measurements from different time points at the same dose were pooled to generate regression curves. Half-maximal inhibitory concentrations (IC₅₀) were determined using a three-parameter logistic regression model. (D) Correlation between transcriptional and translational repression. Data were pooled from 1 and 1.5 nM transfections across days 3, 7, and 14. Error bars: SD. Statistical significance was determined by one-way ANOVA followed by Tukey’s multiple comparisons test, with *P*-values indicated.

**Figure S4.**
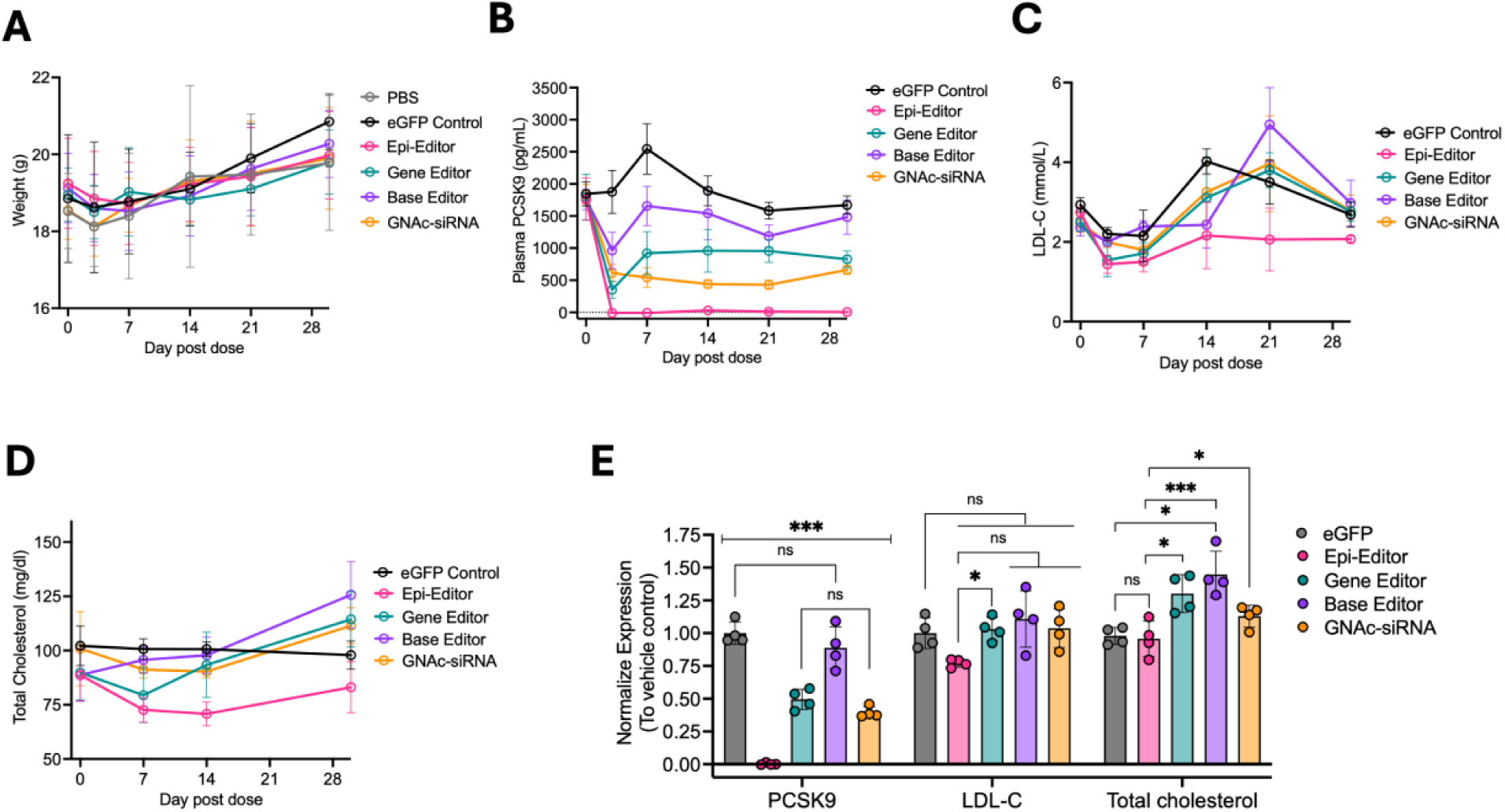
Safety and efficacy of LUNAR^®^–delivered gene modifiers *in vivo*. (A) Body weights of 8–12-week-old female C57BL/6N mice following intravenous administration of 1 mg/kg RNA formulated in LUNAR^®^. (B–D) Plasma levels of PCSK9 protein (B), low-density lipoprotein cholesterol (LDL-C) (C), and total cholesterol (D) measured at indicated time points post-dosing. (E) Comparison of therapeutic efficacy of different gene-modifying modalities on plasma PCSK9, LDL-C, and total cholesterol levels at day 30 post-dose. Statistical significance was determined by one-way ANOVA with Tukey’s multiple comparisons test (\**P* < 0.05, \*\*\**P* < 0.001). For PCSK9 comparisons, differences between groups not specifically indicated are all statistically significant (*P* < 0.001). Error bars: SD.

**Figure S5.**
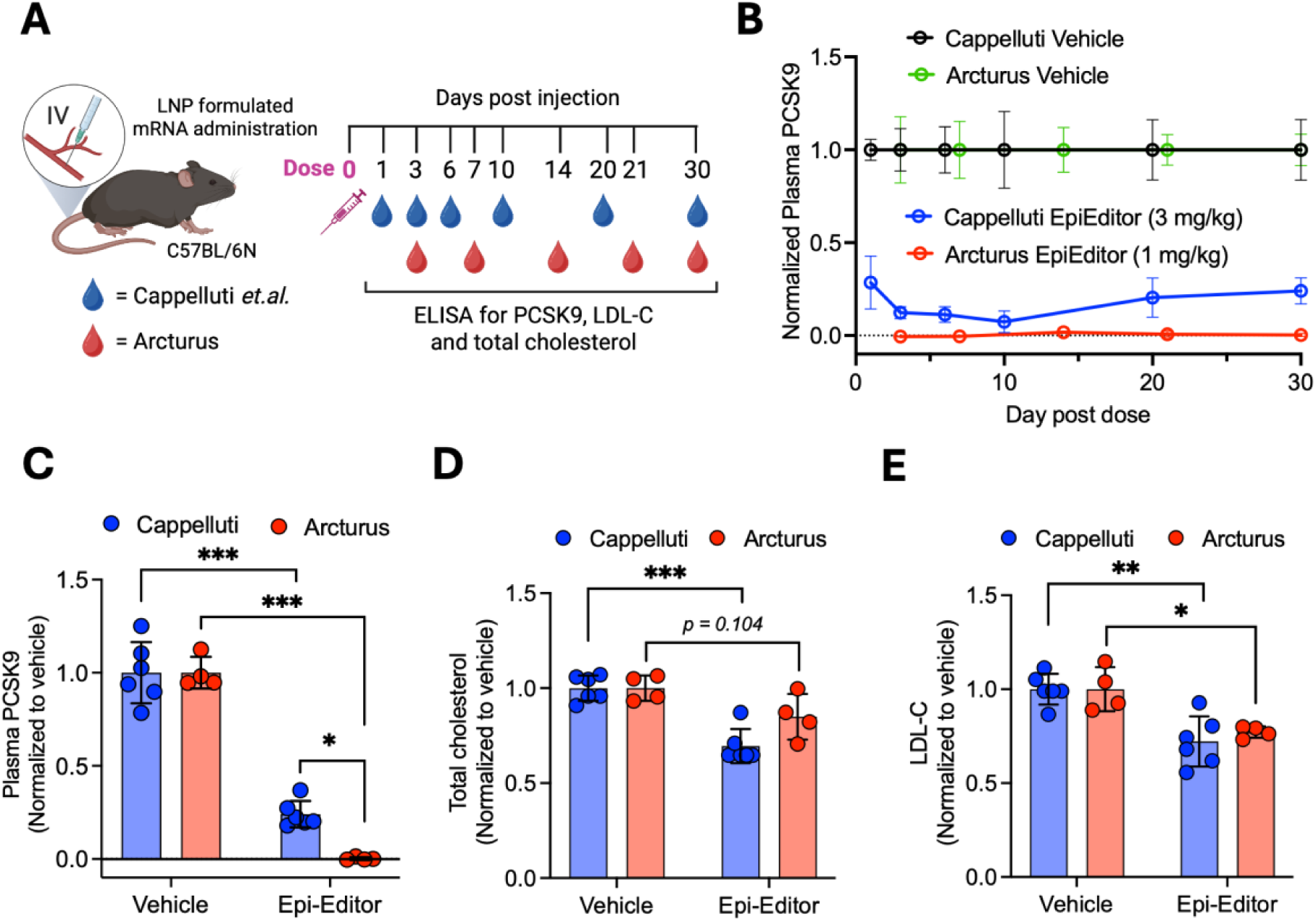
LUNAR^®^ formulation enhances *Pcsk9* suppression even at lower doses *in vivo*. (A) Experimental design comparing the efficacy of epi-editor delivery between the Capelluti et al. study and Arcturus Therapeutics. Plasma was collected at the indicated time points (dots colored by time). (B) Plasma PCSK9 levels measured at each time point post-dosing. Data were normalized to the respective vehicle controls (PBS-treated mice for Capelluti et al. and LUNAR^®^-formulated eGFP mRNA for Arcturus data). (C–E) Comparison of plasma PCSK9 (C), LDL-C (D), and total cholesterol (E) at day 30 post-dosing. Values were normalized to the respective vehicle controls. Error bars: SD. Statistical significance was determined by one-way ANOVA followed by Tukey’s multiple comparisons test (\**P* < 0.05, \*\**P* < 0.01, \*\*\**P* < 0.001). Capelluti’s data were adapted from provided data in Fig. 5B-D and Arcturus data were from the current study.

**Figure S6.**
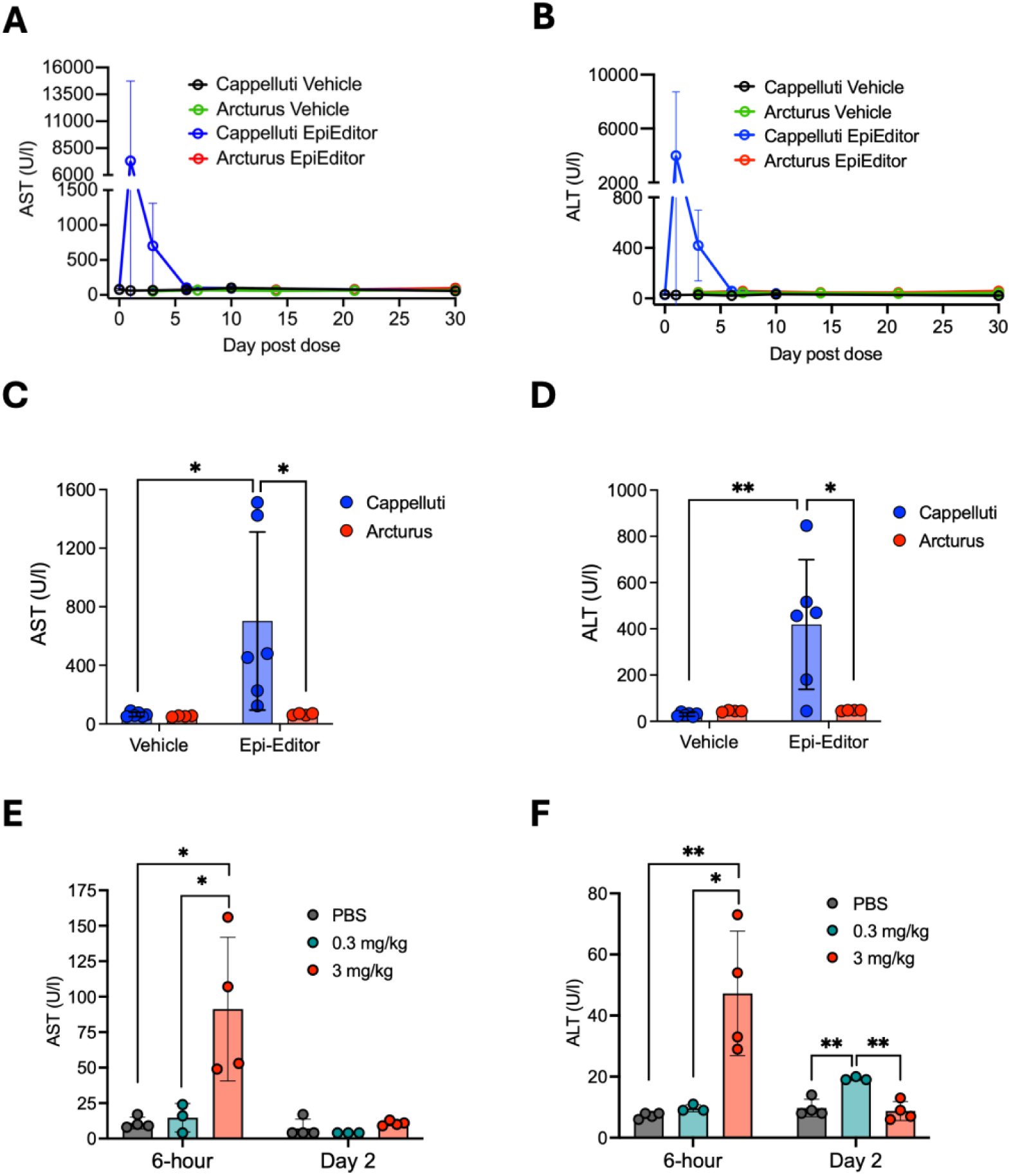
LUNAR^®^ formulation exhibits reduced hepatotoxicity *in vivo*. (A–B) Time-course of serum aspartate aminotransferase (AST) and alanine aminotransferase (ALT) in mice dosed with LNP-formulated mRNA encoding epi-editors or vehicle controls. Vehicle controls were PBS-treated mice in Cappelluti et al. and LUNAR^®^ eGFP mRNA in the Arcturus study. Note that plasma collection time points differed between the two studies. Capelluti’s data were adapted from provided data Fig. 5A and Arcturus data were from the current study. (C–D) Comparison of AST and ALT levels at day 3 post-dosing. Data are derived from panels A and B. Differences between groups are not statistically significant unless indicated. (E–F) Plasma AST and ALT levels in C57BL/6N mice administered human EPO mRNA formulated with LUNAR^®^ at 0.3 or 3 mg/kg, measured at 6 hours and 2 days post-dosing. Data were not part of the current study. Statistical significance was determined by one-way ANOVA followed by Tukey’s multiple comparisons test (\**P* < 0.05, \*\**P* < 0.01). Error bars: SD.

## Supplemental information

### ORF sequence of constructs

#### DNMT-KRAB (Epi-Editor)

MYPYDVPDYASPKKKRKVNHDQEFDPPKVYPPVPAEKRKPIRVLSLFDGIATGLLVLKDL GIQVDRYIASEVCEDSITVGMVRHQGKIMYVGDVRSVTQKHIQEWGPFDLVIGGSPCND LSIVNPARKGLYEGTGRLFFEFYRLLHDARPKEGDDRPFFWLFENVVAMGVSDKRDISRF LESNPVMIDAKEVSAAHRARYFWGNLPGMNRPLASTVNDKLELQECLEHGRIAKFSKV RTITTRSNSIKQGKDQHFPVFMNEKEDILWCTEMERVFGFPVHYTDVSNMSRLARQRLL GRSWSVPVIRHLFAPLKEYFACVSSGNSNANSRGPSFSSGLVPLSLRGSHMAAIPALDPEA EPSMDVILVGSSELSSSVSPGTGRDLIAYEVKANQRNIEDICICCGSLQVHTQHPLFEGGIC APCKDKFLDALFLYDDDGYQSYCSICCSGETLLICGNPDCTRCYCFECVDSLVGPGTSGK VHAMSNWVCYLCLPSSRSGLLQRRRKWRSQLKAFYDRESENPLEMFETVPVWRRQPV RVLSLFEDIKKELTSLGFLESGSDPGQLKHVVDVTDTVRKDVEEWGPFDLVYGATPPLGH TCDRPPSWYLFQFHRLLQYARPKPGSPRPFFWMFVDNLVLNKEDLDVASRFLEMEPVTIP DVHGGSLQNAVRVWSNIPAIRSRHWALVSEEELSLLAQNKQSSKLAAKWPTKLVKNCFL PLREYFKYFSTELTSSLGGPSSGAPPPSGGSPAGSPTSTEEGTSESATPESGPGTSTEPSEGS APGSPAGSPTSTEEGTSTEPSEGSAPGTSTEPSELEGIHGVPAAMAERPFQCRICMRNFSRS DTLSEHIRTHTGEKPFACDICGRKFAHRRSRWGHTKIHTGSQKPFQCRICMRNFSRSAHLS RHIRTHTGEKPFACDICGRKFATSGHLSRHTKIHTGSQKPFQCRICMRNFSERGTLARHIR THTGEKPFACDICGRKFAQSSDLRRHTKIHLRQKDAARGSKLPKKKRKVGSSGSETPGTS ESATPESRTLVTFKDVFVDFTREEWKLLDTAQQIVYRNVMLENYKNLVSLGYQLTKPDVI LRLEKGEEPWLV

#### DNMT

MYPYDVPDYASPKKKRKVNHDQEFDPPKVYPPVPAEKRKPIRVLSLFDGIATGLLVLKDL GIQVDRYIASEVCEDSITVGMVRHQGKIMYVGDVRSVTQKHIQEWGPFDLVIGGSPCND LSIVNPARKGLYEGTGRLFFEFYRLLHDARPKEGDDRPFFWLFENVVAMGVSDKRDISRF LESNPVMIDAKEVSAAHRARYFWGNLPGMNRPLASTVNDKLELQECLEHGRIAKFSKV RTITTRSNSIKQGKDQHFPVFMNEKEDILWCTEMERVFGFPVHYTDVSNMSRLARQRLL GRSWSVPVIRHLFAPLKEYFACVSSGNSNANSRGPSFSSGLVPLSLRGSHMAAIPALDPEA EPSMDVILVGSSELSSSVSPGTGRDLIAYEVKANQRNIEDICICCGSLQVHTQHPLFEGGIC APCKDKFLDALFLYDDDGYQSYCSICCSGETLLICGNPDCTRCYCFECVDSLVGPGTSGK VHAMSNWVCYLCLPSSRSGLLQRRRKWRSQLKAFYDRESENPLEMFETVPVWRRQPV RVLSLFEDIKKELTSLGFLESGSDPGQLKHVVDVTDTVRKDVEEWGPFDLVYGATPPLGH TCDRPPSWYLFQFHRLLQYARPKPGSPRPFFWMFVDNLVLNKEDLDVASRFLEMEPVTIP DVHGGSLQNAVRVWSNIPAIRSRHWALVSEEELSLLAQNKQSSKLAAKWPTKLVKNCFL PLREYFKYFSTELTSSLGGPSSGAPPPSGGSPAGSPTSTEEGTSESATPESGPGTSTEPSEGS APGSPAGSPTSTEEGTSTEPSEGSAPGTSTEPSELEGIHGVPAAMAERPFQCRICMRNFSRS DTLSEHIRTHTGEKPFACDICGRKFAHRRSRWGHTKIHTGSQKPFQCRICMRNFSRSAHLS RHIRTHTGEKPFACDICGRKFATSGHLSRHTKIHTGSQKPFQCRICMRNFSERGTLARHIR THTGEKPFACDICGRKFAQSSDLRRHTKIHLRQKDAARGSKLPKKKRKV

#### KRAB

MYPYDVPDYASPKKKRKVGIHGVPAAMAERPFQCRICMRNFSRSDTLSEHIRTHTGEKP FACDICGRKFAHRRSRWGHTKIHTGSQKPFQCRICMRNFSRSAHLSRHIRTHTGEKPFAC DICGRKFATSGHLSRHTKIHTGSQKPFQCRICMRNFSERGTLARHIRTHTGEKPFACDICG RKFAQSSDLRRHTKIHLRQKDAARGSKLPKKKRKVGSSGSETPGTSESATPESRTLVTFK DVFVDFTREEWKLLDTAQQIVYRNVMLENYKNLVSLGYQLTKPDVILRLEKGEEPWLV

### (-) ZFN Control

MYPYDVPDYASPKKKRKVNHDQEFDPPKVYPPVPAEKRKPIRVLSLFDGIATGLLVLKDL GIQVDRYIASEVCEDSITVGMVRHQGKIMYVGDVRSVTQKHIQEWGPFDLVIGGSPCND LSIVNPARKGLYEGTGRLFFEFYRLLHDARPKEGDDRPFFWLFENVVAMGVSDKRDISRF LESNPVMIDAKEVSAAHRARYFWGNLPGMNRPLASTVNDKLELQECLEHGRIAKFSKV RTITTRSNSIKQGKDQHFPVFMNEKEDILWCTEMERVFGFPVHYTDVSNMSRLARQRLL GRSWSVPVIRHLFAPLKEYFACVSSGNSNANSRGPSFSSGLVPLSLRGSHMAAIPALDPEA EPSMDVILVGSSELSSSVSPGTGRDLIAYEVKANQRNIEDICICCGSLQVHTQHPLFEGGIC APCKDKFLDALFLYDDDGYQSYCSICCSGETLLICGNPDCTRCYCFECVDSLVGPGTSGK VHAMSNWVCYLCLPSSRSGLLQRRRKWRSQLKAFYDRESENPLEMFETVPVWRRQPV RVLSLFEDIKKELTSLGFLESGSDPGQLKHVVDVTDTVRKDVEEWGPFDLVYGATPPLGH TCDRPPSWYLFQFHRLLQYARPKPGSPRPFFWMFVDNLVLNKEDLDVASRFLEMEPVTIP DVHGGSLQNAVRVWSNIPAIRSRHWALVSEEELSLLAQNKQSSKLAAKWPTKLVKNCFL PLREYFKYFSTELTSSLGGPSSGAPPPSGGSPAGSPTSTEEGTSESATPESGPGTSTEPSEGS APGSPAGSPTSTEEGTSTEPSEGSAPGTSTEPSELEGIHGSGKLPKKKRKVGSSGSETPGTS ESATPESRTLVTFKDVFVDFTREEWKLLDTAQQIVYRNVMLENYKNLVSLGYQLTKPDVI LRLEKGEEPWLV

#### ZFN Control

MYPYDVPDYASPKKKRKVGIHGVPAAMAERPFQCRICMRNFSRSDTLSEHIRTHTGEKP FACDICGRKFAHRRSRWGHTKIHTGSQKPFQCRICMRNFSRSAHLSRHIRTHTGEKPFAC DICGRKFATSGHLSRHTKIHTGSQKPFQCRICMRNFSERGTLARHIRTHTGEKPFACDICG RKFAQSSDLRRHTKIHLRQKDAARGSKLPKKKRKV

#### Gene editor

MGPKKKRKVGGSPKKKRKVGGSPKKKRKVGGSPKKKRKVGGGSMDKKYSIGLDIGTN SVGWAVITDEYKVPSKKFKVLGNTDRHSIKKNLIGALLFDSGETAEATRLKRTARRRYTR RKNRICYLQEIFSNEMAKVDDSFFHRLEESFLVEEDKKHERHPIFGNIVDEVAYHEKYPTI YHLRKKLVDSTDKADLRLIYLALAHMIKFRGHFLIEGDLNPDNSDVDKLFIQLVQTYNQ LFEENPINASGVDAKAILSARLSKSRRLENLIAQLPGEKKNGLFGNLIALSLGLTPNFKSN FDLAEDAKLQLSKDTYDDDLDNLLAQIGDQYADLFLAAKNLSDAILLSDILRVNTEITKA PLSASMIKRYDEHHQDLTLLKALVRQQLPEKYKEIFFDQSKNGYAGYIDGGASQEEFYKF IKPILEKMDGTEELLVKLNREDLLRKQRTFDNGSIPHQIHLGELHAILRRQEDFYPFLKDN REKIEKILTFRIPYYVGPLARGNSRFAWMTRKSEETITPWNFEEVVDKGASAQSFIERMTA FDKNLPNEKVLPKHSLLYEYFTVYNELTKVKYVTEGMRKPAFLSGEQKKAIVDLLFKTN RKVTVKQLKEDYFKKIECFDSVEISGVEDRFNASLGTYHDLLKIIKDKDFLDNEENEDILE DIVLTLTLFEDREMIEERLKTYAHLFDDKVMKQLKRRRYTGWGALSRKLINGIRDKQSG KTILDFLKSDGFANRNFMALIHDDSLTFKEDIQKAQVSGQGDSLHEHIANLAGSPAIKKGI LQTVKVVDELVKVMGRHKPENIVIEMARENQTTQKGQKNSRERMKRIEEGIKELGSQIL KEHPVENTQLQNEKLYLYYLQNGRDMYVDQELDINRLSDYDVDHIVPQSFLKDDSIDNK VLTRSDKNRGKSDNVPSEEVVKKMKNYWRQLLNAKLITQRKFDNLTKAERGGLSELDK AGFIKRQLVETRAITKHVAQILDSRMNTKYDENDKLIREVKVITLKSKLVSDFRKDFQFY KVREINNYHHAHDAYLNAVVGTALIKKYPKLESEFVYGDYKVYDVRKMIAKSEQEIGK ATAKYFFYSNIMNFFKTEITLANGEIRKRPLIETNGETGEIVWDKGRDFATVRKVLSMPQ VNIVKKTEVQTGGFSKESILPKRNSDKLIARKKDWDPKKYGGFDSPTVAYSVLVVAKVE KGKSKKLKSVKELLGITIMERSSFEKNPIDFLEAKGYKEVKKDLIIKLPKYSLFELENGRK RMLASAGELQKGNELALPSKYVNFLYLASHYEKLKGSPEDNEQKQLFVEQHKHYLDEII EQISEFSKRVILADANLDKVLSAYNKHRDKPIREQAENIIHLFTLTNLGAPAAFKYFDTTID RKRYTSTKEVLDATLIHQSITGLYETRIDLSQLGGDGGGSPKKKRKV

#### Base editor

MGPKKKRKVGGSPKKKRKVGGSPKKKRKVGGSPKKKRKVGGGSMDKKYSIGLAIGTN SVGWAVITDEYKVPSKKFKVLGNTDRHSIKKNLIGALLFDSGETAEATRLKRTARRRYTR RKNRICYLQEIFSNEMAKVDDSFFHRLEESFLVEEDKKHERHPIFGNIVDEVAYHEKYPTI YHLRKKLVDSTDKADLRLIYLALAHMIKFRGHFLIEGDLNPDNSDVDKLFIQLVQTYNQ LFEENPINASGVDAKAILSARLSKSRRLENLIAQLPGEKKNGLFGNLIALSLGLTPNFKSN FDLAEDAKLQLSKDTYDDDLDNLLAQIGDQYADLFLAAKNLSDAILLSDILRVNTEITKA PLSASMIKRYDEHHQDLTLLKALVRQQLPEKYKEIFFDQSKNGYAGYIDGGASQEEFYKF IKPILEKMDGTEELLVKLNREDLLRKQRTFDNGSIPHQIHLGELHAILRRQEDFYPFLKDN REKIEKILTFRIPYYVGPLARGNSRFAWMTRKSEETITPWNFEEVVDKGASAQSFIERMTA FDKNLPNEKVLPKHSLLYEYFTVYNELTKVKYVTEGMRKPAFLSGEQKKAIVDLLFKTN RKVTVKQLKEDYFKKIECFDSVEISGVEDRFNASLGTYHDLLKIIKDKDFLDNEENEDILE DIVLTLTLFEDREMIEERLKTYAHLFDDKVMKQLKRRRYTGWGALSRKLINGIRDKQSG KTILDFLKSDGFANRNFMALIHDDSLTFKEDIQKAQVSGQGDSLHEHIANLAGSPAIKKGI LQTVKVVDELVKVMGRHKPENIVIEMARENQTTQKGQKNSRERMKRIEEGIKELGSQIL KEHPVENTQLQNEKLYLYYLQNGRDMYVDQELDINRLSDYDVDHIVPQSFLKDDSIDNK VLTRSDKNRGKSDNVPSEEVVKKMKNYWRQLLNAKLITQRKFDNLTKAERGGLSELDK AGFIKRQLVETRAITKHVAQILDSRMNTKYDENDKLIREVKVITLKSKLVSDFRKDFQFY KVREINNYHHAHDAYLNAVVGTALIKKYPKLESEFVYGDYKVYDVRKMIAKSEQEIGK ATAKYFFYSNIMNFFKTEITLANGEIRKRPLIETNGETGEIVWDKGRDFATVRKVLSMPQ VNIVKKTEVQTGGFSKESILPKRNSDKLIARKKDWDPKKYGGFDSPTVAYSVLVVAKVE KGKSKKLKSVKELLGITIMERSSFEKNPIDFLEAKGYKEVKKDLIIKLPKYSLFELENGRK RMLASAGELQKGNELALPSKYVNFLYLASHYEKLKGSPEDNEQKQLFVEQHKHYLDEII EQISEFSKRVILADANLDKVLSAYNKHRDKPIREQAENIIHLFTLTNLGAPAAFKYFDTTID RKRYTSTKEVLDATLIHQSITGLYETRIDLSQLGGDGSPKKKRKVGGGGSGGGGSAEYV RALFDFNGNDEEDLPFKKGDILRIRDKPEEQWWNAEDSEGKRGMIPVPYVEKYSGDYK DHDGDYKDHDIDYKDDDDKSRMTDAEYVRIHEKLDIYTFKKQFFNNKKSVSHRCYVLF ELKRRGERRACFWGYAVNKPQSGTERGIHAEIFSIRKVEEYLRDNPGQFTINWYSSWSPC ADCAEKILEWYNQELRGNGHTLKIWACKLYYEKNARNQIGLWNLRDNGVGLNVMVSE HYQCCRKIFIQSSHNQLNENRWLEKTLKRAEKRRSELSIMIQVKILHTTKSPAVGTPKKK RKVGTMTNLSDIIEKETGKQLVIQESILMLPEEVEEVIGNKPESDILVHTAYDESTDENVM LLTSDAPEYKPWALVIQDSNGENKIKML

#### eGFP mock control

MVSKGEELFTGVVPILVELDGDVNGHKFSVSGEGEGDATYGKLTLKFICTTGKLPVPWP TLVTTLTYGVQCFSRYPDHMKQHDFFKSAMPEGYVQERTIFFKDDGNYKTRAEVKFEG DTLVNRIELKGIDFKEDGNILGHKLEYNYNSHNVYIMADKQKNGIKVNFKIRHNIEDGSV QLADHYQQNTPIGDGPVLLPDNHYLSTQSALSKDPNEKRDHMVLLEFVTAAGITLGMD ELYK

#### Firefly luciferase

MEDAKNIKKGPAPFYPLEDGTAGEQLHKAMKRYALVPGTIAFTDAHIEVDITYAEYFEMS VRLAEAMKRYGLNTNHRIVVCSENSLQFFMPVLGALFIGVAVAPANDIYNERELLNSMGI SQPTVVFVSKKGLQKILNVQKKLPIIQKIIIMDSKTDYQGFQSMYTFVTSHLPPGFNEYDF VPESFDRDKTIALIMNSSGSTGLPKGVALPHRTACVRFSHARDPIFGNQIIPDTAILSVVPF HHGFGMFTTLGYLICGFRVVLMYRFEEELFLRSLQDYKIQSALLVPTLFSFFAKSTLIDKY DLSNLHEIASGGAPLSKEVGEAVAKRFHLPGIRQGYGLTETTSAILITPEGDDKPGAVGKV VPFFEAKVVDLDTGKTLGVNQRGELCVRGPMIMSGYVNNPEATNALIDKDGWLHSGDI AYWDEDEHFFIVDRLKSLIKYKGYQVAPAELESILLQHPNIFDAGVAGLPDDDAGELPAA VVVLEHGKTMTEKEIVDYVASQVTTAKKLRGGVVFVDEVPKGLTGKLDARKIREILIKA KKGGKIAV

#### GalNAc-siRNA

Sense: 5’-AGGCCUGGAGUUUAUUCGGAA-3’ Antisense: 5’-UUCCGAAUAAACUCCAGGCCUAU-3’

#### sgRNA sequences

sgRNA1 target sequence: 5’-CCGCAGCCACGCAGAGCAGT-3’

sgRNA2 target sequence: 5’-ACCGCAGCCACGCAGAGCAG-3’

## References

1. Lambert, G., Sjouke, B., Choque, B., Kastelein, J.J., and Hovingh, G.K. (2012). The PCSK9 decade: thematic review series: new lipid and lipoprotein targets for the treatment of cardiometabolic diseases. Journal of lipid research 53, 2515–2524.

2. Surdo, P.L., Bottomley, M.J., Calzetta, A., Settembre, E.C., Cirillo, A., Pandit, S., Ni, Y.G., Hubbard, B., Sitlani, A., and Carfí, A. (2011). Mechanistic implications for LDL receptor degradation from the PCSK9/LDLR structure at neutral pH. EMBO reports 12, 1300–1305.

3. Page, M.M., and Watts, G.F. (2016). PCSK9 inhibitors–mechanisms of action. Australian prescriber 39, 164.

4. Corsini, A., Ginsberg, H.N., and Chapman, M.J. (2025). Therapeutic PCSK9 targeting: Inside versus outside the hepatocyte? Pharmacology & Therapeutics, 108812.

5. Hobbs, H.H., Cohen, J.C., and Horton, J.D. (2024). PCSK9: from nature’s loss to patient’s gain. Circulation 149, 171–173.

6. Cohen, J.C., Boerwinkle, E., Mosley Jr, T.H., and Hobbs, H.H. (2006). Sequence variations in PCSK9, low LDL, and protection against coronary heart disease. New England Journal of Medicine 354, 1264–1272.

7. Kent, S.T., Rosenson, R.S., Avery, C.L., Chen, Y.-D.I., Correa, A., Cummings, S.R., Cupples, L.A., Cushman, M., Evans, D.S., and Gudnason, V. (2017). PCSK9 loss-of-function variants, low-density lipoprotein cholesterol, and risk of coronary heart disease and stroke: data from 9 studies of blacks and whites. Circulation: cardiovascular genetics 10, e001632.

8. Langsted, A., Nordestgaard, B.G., Benn, M., Tybjærg-Hansen, A., and Kamstrup, P.R. (2016). PCSK9 R46L loss-of-function mutation reduces lipoprotein (a), LDL cholesterol, and risk of aortic valve stenosis. The Journal of Clinical Endocrinology & Metabolism 101, 3281–3287.

9. Sabatine, M.S., Giugliano, R.P., Keech, A.C., Honarpour, N., Wiviott, S.D., Murphy, S.A., Kuder, J.F., Wang, H., Liu, T., and Wasserman, S.M. (2017). Evolocumab and clinical outcomes in patients with cardiovascular disease. New England journal of medicine 376, 1713–1722.

10. Stein, E.A., Mellis, S., Yancopoulos, G.D., Stahl, N., Logan, D., Smith, W.B., Lisbon, E., Gutierrez, M., Webb, C., and Wu, R. (2012). Effect of a monoclonal antibody to PCSK9 on LDL cholesterol. New England Journal of Medicine 366, 1108–1118.

11. Hassan, M. (2015). OSLER and ODYSSEY LONG TERM: PCSK9 inhibitors on the right track of reducing cardiovascular events. Global Cardiology Science and Practice 2015, 20.

12. Raal, F.J., Kallend, D., Ray, K.K., Turner, T., Koenig, W., Wright, R.S., Wijngaard, P.L., Curcio, D., Jaros, M.J., and Leiter, L.A. (2020). Inclisiran for the treatment of heterozygous familial hypercholesterolemia. New England Journal of Medicine 382, 1520–1530.

13. Fernández-Ruiz, I. (2020). Twice-yearly inclisiran injections halve LDL-cholesterol levels. Nature Reviews Cardiology 17, 321–322.

14. Ray, K.K., Troquay, R.P., Visseren, F.L., Leiter, L.A., Wright, R.S., Vikarunnessa, S., Talloczy, Z., Zang, X., Maheux, P., and Lesogor, A. (2023). Long-term efficacy and safety of inclisiran in patients with high cardiovascular risk and elevated LDL cholesterol (ORION-3): results from the 4-year open-label extension of the ORION-1 trial. The Lancet Diabetes & Endocrinology 11, 109–119.

15. Wang, X., Raghavan, A., Chen, T., Qiao, L., Zhang, Y., Ding, Q., and Musunuru, K. (2016). CRISPR-Cas9 targeting of PCSK9 in human hepatocytes in vivo—brief report. Arteriosclerosis, thrombosis, and vascular biology 36, 783–786.

16. Jinek, M., Chylinski, K., Fonfara, I., Hauer, M., Doudna, J.A., and Charpentier, E. (2012). A programmable dual-RNA–guided DNA endonuclease in adaptive bacterial immunity. science 337, 816–821.

17. Jinek, M., East, A., Cheng, A., Lin, S., Ma, E., and Doudna, J. (2013). RNA-programmed genome editing in human cells. elife 2, e00471.

18. Nishimasu, H., Shi, X., Ishiguro, S., Gao, L., Hirano, S., Okazaki, S., Noda, T., Abudayyeh, O.O., Gootenberg, J.S., and Mori, H. (2018). Engineered CRISPR-Cas9 nuclease with expanded targeting space. Science 361, 1259–1262.

19. Kleinstiver, B.P., Pattanayak, V., Prew, M.S., Tsai, S.Q., Nguyen, N.T., Zheng, Z., and Joung, J.K. (2016). High-fidelity CRISPR–Cas9 nucleases with no detectable genome-wide off-target effects. Nature 529, 490–495.

20. Wang, D., Zhang, C., Wang, B., Li, B., Wang, Q., Liu, D., Wang, H., Zhou, Y., Shi, L., and Lan, F. (2019). Optimized CRISPR guide RNA design for two high-fidelity Cas9 variants by deep learning. Nature communications 10, 4284.

21. Kuscu, C., Arslan, S., Singh, R., Thorpe, J., and Adli, M. (2014). Genome-wide analysis reveals characteristics of off-target sites bound by the Cas9 endonuclease. Nature biotechnology 32, 677–683.

22. Kosicki, M., Tomberg, K., and Bradley, A. (2018). Repair of double-strand breaks induced by CRISPR–Cas9 leads to large deletions and complex rearrangements. Nature biotechnology 36, 765–771.

23. Komor, A.C., Kim, Y.B., Packer, M.S., Zuris, J.A., and Liu, D.R. (2016). Programmable editing of a target base in genomic DNA without double-stranded DNA cleavage. Nature 533, 420–424.

24. Gaudelli, N.M., Komor, A.C., Rees, H.A., Packer, M.S., Badran, A.H., Bryson, D.I., and Liu, D.R. (2017). Programmable base editing of A• T to G• C in genomic DNA without DNA cleavage. Nature 551, 464–471.

25. Gehrke, J.M., Cervantes, O., Clement, M.K., Wu, Y., Zeng, J., Bauer, D.E., Pinello, L., and Joung, J.K. (2018). An APOBEC3A-Cas9 base editor with minimized bystander and off-target activities. Nature biotechnology 36, 977–982.

26. Musunuru, K., Grandinette, S.A., Wang, X., Hudson, T.R., Briseno, K., Berry, A.M., Hacker, J.L., Hsu, A., Silverstein, R.A., and Hille, L.T. (2025). Patient-specific in vivo gene editing to treat a rare genetic disease. New England Journal of Medicine 392, 2235–2243.

27. Cappelluti, M.A., Mollica Poeta, V., Valsoni, S., Quarato, P., Merlin, S., Merelli, I., and Lombardo, A. (2024). Durable and efficient gene silencing in vivo by hit-and-run epigenome editing. Nature 627, 416–423.

28. Xu, D., Besselink, S., Ramadoss, G.N., Dierks, P.H., Lubin, J.P., Pattali, R.K., Brim, J.I., Christenson, A.E., Colias, P.J., and Ornelas, I.J. (2025). Programmable epigenome editing by transient delivery of CRISPR epigenome editor ribonucleoproteins. Nature Communications 16, 7948.

29. Nuñez, J.K., Chen, J., Pommier, G.C., Cogan, J.Z., Replogle, J.M., Adriaens, C., Ramadoss, G.N., Shi, Q., Hung, K.L., and Samelson, A.J. (2021). Genome-wide programmable transcriptional memory by CRISPR-based epigenome editing. Cell 184, 2503–2519. e2517.

30. Thakore, P.I., D’ippolito, A.M., Song, L., Safi, A., Shivakumar, N.K., Kabadi, A.M., Reddy, T.E., Crawford, G.E., and Gersbach, C.A. (2015). Highly specific epigenome editing by CRISPR-Cas9 repressors for silencing of distal regulatory elements. Nature methods 12, 1143–1149.

31. Tremblay, F., Xiong, Q., Shah, S.S., Ko, C.-W., Kelly, K., Morrison, M.S., Giancarlo, C., Ramirez, R.N., Hildebrand, E.M., and Voytek, S.B. (2025). A potent epigenetic editor targeting human PCSK9 for durable reduction of low-density lipoprotein cholesterol levels. Nature medicine 31, 1329–1338.

32. Bernstein, D.L., Le Lay, J.E., Ruano, E.G., and Kaestner, K.H. (2015). TALE-mediated epigenetic suppression of CDKN2A increases replication in human fibroblasts. The Journal of clinical investigation 125, 1998–2006.

33. Mendenhall, E.M., Williamson, K.E., Reyon, D., Zou, J.Y., Ram, O., Joung, J.K., and Bernstein, B.E. (2013). Locus-specific editing of histone modifications at endogenous enhancers. Nature biotechnology 31, 1133–1136.

34. Schulze, R.J., Schott, M.B., Casey, C.A., Tuma, P.L., and McNiven, M.A. (2019). The cell biology of the hepatocyte: A membrane trafficking machine. Journal of Cell Biology 218, 2096–2112.

35. Issa, S.S., Shaimardanova, A.A., Solovyeva, V.V., and Rizvanov, A.A. (2023). Various AAV serotypes and their applications in gene therapy: an overview. Cells 12, 785.

36. Li, Q., Su, J., Liu, Y., Jin, X., Zhong, X., Mo, L., Wang, Q., Deng, H., and Yang, Y. (2021). In vivo PCSK9 gene editing using an all-in-one self-cleavage AAV-CRISPR system. Molecular Therapy Methods & Clinical Development 20, 652–659.

37. Garcia, D.A., Pierre, A.F., Quirino, L., Acharya, G., Vasudevan, A., Pei, Y., Chung, E., Chang, J.Y., Lee, S., and Endow, M. (2025). Lipid nanoparticle delivery of TALEN mRNA targeting LPA causes gene disruption and plasma lipoprotein (a) reduction in transgenic mice. Molecular Therapy 33, 90–103.

38. Ramaswamy, S., Tonnu, N., Tachikawa, K., Limphong, P., Vega, J.B., Karmali, P.P., Chivukula, P., and Verma, I.M. (2017). Systemic delivery of factor IX messenger RNA for protein replacement therapy. Proceedings of the National Academy of Sciences 114, E1941–E1950.

39. Zhang, T., Yin, H., Li, Y., Yang, H., Ge, K., Zhang, J., Yuan, Q., Dai, X., Naeem, A., and Weng, Y. (2024). Optimized lipid nanoparticles (LNPs) for organ-selective nucleic acids delivery in vivo. Iscience 27.

40. Kim, M., Jeong, M., Hur, S., Cho, Y., Park, J., Jung, H., Seo, Y., Woo, H., Nam, K., and Lee, K. (2021). Engineered ionizable lipid nanoparticles for targeted delivery of RNA therapeutics into different types of cells in the liver. Science advances 7, eabf4398.

41. Katzmann, J.L., Cupido, A.J., and Laufs, U. (2022). Gene therapy targeting PCSK9. Metabolites 12, 70.

42. Musunuru, K., Chadwick, A.C., Mizoguchi, T., Garcia, S.P., DeNizio, J.E., Reiss, C.W., Wang, K., Iyer, S., Dutta, C., and Clendaniel, V. (2021). In vivo CRISPR base editing of PCSK9 durably lowers cholesterol in primates. Nature 593, 429–434.

43. Arribere, J.A., and Fire, A.Z. (2018). Nonsense mRNA suppression via nonstop decay. Elife 7, e33292.

44. Brogna, S., and Wen, J. (2009). Nonsense-mediated mRNA decay (NMD) mechanisms. Nature structural & molecular biology 16, 107–113.

45. Ran, F.A., Hsu, P.D., Lin, C.-Y., Gootenberg, J.S., Konermann, S., Trevino, A.E., Scott, D.A., Inoue, A., Matoba, S., and Zhang, Y. (2013). Double nicking by RNA-guided CRISPR Cas9 for enhanced genome editing specificity. Cell 154, 1380–1389.

46. Tan, J., Zhang, F., Karcher, D., and Bock, R. (2019). Engineering of high-precision base editors for site-specific single nucleotide replacement. Nat. Commun. 10, 439.

47. Nishida, K., Arazoe, T., Yachie, N., Banno, S., Kakimoto, M., Tabata, M., Mochizuki, M., Miyabe, A., Araki, M., and Hara, K.Y. (2016). Targeted nucleotide editing using hybrid prokaryotic and vertebrate adaptive immune systems. Science 353, aaf8729.

48. Doman, J.L., Raguram, A., Newby, G.A., and Liu, D.R. (2020). Evaluation and minimization of Cas9-independent off-target DNA editing by cytosine base editors. Nature biotechnology 38, 620–628.

49. Hwang, G.-H., Park, J., Lim, K., Kim, S., Yu, J., Yu, E., Kim, S.-T., Eils, R., Kim, J.-S., and Bae, S. (2018). Web-based design and analysis tools for CRISPR base editing. BMC bioinformatics 19, 542.

50. Biba, D., Klink, G., and Bazykin, G.A. (2022). Pairs of mutually compensatory frameshifting mutations contribute to protein evolution. Molecular Biology and Evolution 39, msac031.

51. Groner, A.C., Meylan, S., Ciuffi, A., Zangger, N., Ambrosini, G., Dénervaud, N., Bucher, P., and Trono, D. (2010). KRAB–zinc finger proteins and KAP1 can mediate long-range transcriptional repression through heterochromatin spreading. PLoS genetics 6, e1000869.

52. Fuks, F., Hurd, P.J., Deplus, R., and Kouzarides, T. (2003). The DNA methyltransferases associate with HP1 and the SUV39H1 histone methyltransferase. Nucleic acids research 31, 2305–2312.

53. Morris, M.J., and Monteggia, L.M. (2014). Role of DNA methylation and the DNA methyltransferases in learning and memory. Dialogues in clinical neuroscience 16, 359–371.

54. Suetake, I., Shinozaki, F., Miyagawa, J., Takeshima, H., and Tajima, S. (2004). DNMT3L stimulates the DNA methylation activity of Dnmt3a and Dnmt3b through a direct interaction. Journal of Biological Chemistry 279, 27816–27823.

55. Chedin, F., Lieber, M.R., and Hsieh, C.-L. (2002). The DNA methyltransferase-like protein DNMT3L stimulates de novo methylation by Dnmt3a. Proceedings of the National Academy of Sciences 99, 16916–16921.

56. Siddique, A.N., Nunna, S., Rajavelu, A., Zhang, Y., Jurkowska, R.Z., Reinhardt, R., Rots, M.G., Ragozin, S., Jurkowski, T.P., and Jeltsch, A. (2013). Targeted methylation and gene silencing of VEGF-A in human cells by using a designed Dnmt3a–Dnmt3L single-chain fusion protein with increased DNA methylation activity. Journal of molecular biology 425, 479–491.

57. Jeltsch, A., Broche, J., and Bashtrykov, P. (2018). Molecular processes connecting DNA methylation patterns with DNA methyltransferases and histone modifications in mammalian genomes. Genes 9, 566.

58. Mudla, A., Jiang, Y., Arimoto, K.-i., Xu, B., Rajesh, A., Ryan, A.P., Wang, W., Daugherty, M.D., Zhang, D.-E., and Hao, N. (2020). Cell-cycle-gated feedback control mediates desensitization to interferon stimulation. Elife 9, e58825.

59. Pękowska, A. (2025). The dimmer switch in epigenetics: How DNA methylation encodes gene activity over time. Cell Genomics 5.

60. Sarkies, P., and Sale, J.E. (2012). Propagation of histone marks and epigenetic memory during normal and interrupted DNA replication. Cellular and Molecular Life Sciences 69, 697–716.

61. Wenger, A., Biran, A., Alcaraz, N., Redó-Riveiro, A., Sell, A.C., Krautz, R., Flury, V., Reverón-Gómez, N., Solis-Mezarino, V., and Völker-Albert, M. (2023). Symmetric inheritance of parental histones governs epigenome maintenance and embryonic stem cell identity. Nature genetics 55, 1567–1578.

62. Ozer, J., Ratner, M., Shaw, M., Bailey, W., and Schomaker, S. (2008). The current state of serum biomarkers of hepatotoxicity. Toxicology 245, 194–205.

63. Xia, X.-d., Peng, Z.-s., Gu, H.-m., Wang, M., Wang, G.-q., and Zhang, D.-w. (2021). Regulation of PCSK9 expression and function: mechanisms and therapeutic implications. Frontiers in cardiovascular medicine 8, 764038.

64. Bao, X., Liang, Y., Chang, H., Cai, T., Feng, B., Gordon, K., Zhu, Y., Shi, H., He, Y., and Xie, L. (2024). Targeting proprotein convertase subtilisin/kexin type 9 (PCSK9): from bench to bedside. Signal transduction and targeted therapy 9, 13.

65. Gordon, S.M., Li, H., Zhu, X., Shah, A.S., Lu, L.J., and Davidson, W.S. (2015). A comparison of the mouse and human lipoproteome: suitability of the mouse model for studies of human lipoproteins. Journal of proteome research 14, 2686–2695.

66. Rothgangl, T., Dennis, M.K., Lin, P.J., Oka, R., Witzigmann, D., Villiger, L., Qi, W., Hruzova, M., Kissling, L., and Lenggenhager, D. (2021). In vivo adenine base editing of PCSK9 in macaques reduces LDL cholesterol levels. Nature biotechnology 39, 949–957.

67. Park, S., and Beal, P.A. (2019). Off-target editing by CRISPR-guided DNA base editors. Biochemistry 58, 3727–3734.

68. Kim, Y.B., Komor, A.C., Levy, J.M., Packer, M.S., Zhao, K.T., and Liu, D.R. (2017). Increasing the genome-targeting scope and precision of base editing with engineered Cas9-cytidine deaminase fusions. Nature biotechnology 35, 371–376.

69. Sontag, T.J., Chellan, B., Getz, G.S., and Reardon, C.A. (2013). Differing rates of cholesterol absorption among inbred mouse strains yield differing levels of HDL-cholesterol. Journal of Lipid Research 54, 2515–2524.

70. Berger, J.-M., Valdes, A.L., Gromada, J., Anderson, N., and Horton, J.D. (2017). Inhibition of PCSK9 does not improve lipopolysaccharide-induced mortality in mice. Journal of lipid research 58, 1661–1669.

71. Rashid, S., Curtis, D.E., Garuti, R., Anderson, N.N., Bashmakov, Y., Ho, Y., Hammer, R.E., Moon, Y.-A., and Horton, J.D. (2005). Decreased plasma cholesterol and hypersensitivity to statins in mice lacking Pcsk9. Proceedings of the National Academy of Sciences 102, 5374–5379.

72. Grefhorst, A., McNutt, M.C., Lagace, T.A., and Horton, J.D. (2008). Plasma PCSK9 preferentially reduces liver LDL receptors in mice. Journal of lipid research 49, 1303–1311.

73. Schulz, R., and Schlüter, K.-D. (2017). PCSK9 targets important for lipid metabolism. Clinical research in cardiology supplements 12, 2–11.

74. Carreras, A., Pane, L.S., Nitsch, R., Madeyski-Bengtson, K., Porritt, M., Akcakaya, P., Taheri-Ghahfarokhi, A., Ericson, E., Bjursell, M., and Perez-Alcazar, M. (2019). In vivo genome and base editing of a human PCSK9 knock-in hypercholesterolemic mouse model. BMC biology 17, 4.

75. Keeter, W.C., Carter, N.M., Nadler, J.L., and Galkina, E.V. (2022). The AAV-PCSK9 murine model of atherosclerosis and metabolic dysfunction. European Heart Journal Open 2, oeac028.

76. Tóth, M.E., Dukay, B., Hoyk, Z., and Sántha, M. (2020). Cerebrovascular changes and neurodegeneration related to hyperlipidemia: characteristics of the human ApoB-100 transgenic mice. Current Pharmaceutical Design 26, 1486–1494.

77. Frank-Kamenetsky, M., Grefhorst, A., Anderson, N.N., Racie, T.S., Bramlage, B., Akinc, A., Butler, D., Charisse, K., Dorkin, R., and Fan, Y. (2008). Therapeutic RNAi targeting PCSK9 acutely lowers plasma cholesterol in rodents and LDL cholesterol in nonhuman primates. Proceedings of the National Academy of Sciences 105, 11915–11920.

78. Wang, J., Ding, Y., Chong, K., Cui, M., Cao, Z., Tang, C., Tian, Z., Hu, Y., Zhao, Y., and Jiang, S. (2024). Recent advances in lipid nanoparticles and their safety concerns for mRNA delivery. Vaccines 12, 1148.

79. Liu, Y., Huang, Y., He, G., Guo, C., Dong, J., and Wu, L. (2024). Development of mRNA lipid nanoparticles: targeting and therapeutic aspects. International Journal of Molecular Sciences 25, 10166.

80. Yuan, Z., Yan, R., Fu, Z., Wu, T., and Ren, C. (2024). Impact of physicochemical properties on biological effects of lipid nanoparticles: Are they completely safe. Science of The Total Environment 927, 172240.

81. McGill, M.R. (2016). The past and present of serum aminotransferases and the future of liver injury biomarkers. EXCLI journal 15, 817.

82. Perez-Garcia, C.G., Diaz-Trelles, R., Vega, J.B., Bao, Y., Sablad, M., Limphong, P., Chikamatsu, S., Yu, H., Taylor, W., and Karmali, P.P. (2022). Development of an mRNA replacement therapy for phenylketonuria. Molecular Therapy Nucleic Acids 28, 87–98.

83. De Alwis, R., Gan, E.S., Chen, S., Leong, Y.S., Tan, H.C., Zhang, S.L., Yau, C., Low, J.G., Kalimuddin, S., and Matsuda, D. (2021). A single dose of self-transcribing and replicating RNA-based SARS-CoV-2 vaccine produces protective adaptive immunity in mice. Molecular Therapy 29, 1970–1983.

84. Mukthavaram, R., Sagi, A., Hong, H., Recatto, R., Chikamatsu, S., Leu, A., Shishina, A., Acharya, G., Dam, T., and Soontornniyomkij, B. (2025). Design and Evaluation of Biodegradable Self-Immolative Lipids for RNA Delivery. Advanced Healthcare Materials 14, 2501305.

